# Oncogenic ZMYND11-MBTD1 fusion protein anchors the NuA4/TIP60 histone acetyltransferase complex to the coding region of active gene

**DOI:** 10.1101/2021.03.08.434474

**Authors:** Maëva Devoucoux, Victoire Fort, Gabriel Khelifi, Joshua Xu, Nader Alerasool, Maxime Galloy, Nicholas Wong, Gaëlle Bourriquen, Amélie Fradet-Turcotte, Mikko Taipale, Kristin Hope, Samer M. I. Hussein, Jacques Côté

**Affiliations:** St. Patrick Research Group in Basic Oncology, Laval University Cancer Research Center, Oncology division of CHU de Québec-Université Laval Research Center, Quebec City, QC G1R 3S3, Canada; Department of Biochemistry and Biomedical Sciences, McMaster University, Hamilton, ON L8S 4K1, Canada; Donnelly Centre for Cellular and Biomolecular Research, Department of Molecular Genetics, University of Toronto, Toronto, ON M5S 3E1, Canada; Princess Margaret Cancer Centre, University Health Network, Toronto, ON, M5G 1L7, Canada; Department of Medical Biophysics, University of Toronto, Toronto, ON, M5G 1L7, Canada

**Keywords:** ZMYND11, MBTD1, NuA4, TIP60, AML, translocation, protein fusion, MYC, chromatin, acetylation, transcription, mRNA processing

## Abstract

A chromosomal translocation found in cannibalistic acute myeloid leukemia (AML) leads to an in-frame fusion of the transcription elongation repressor ZMYND11 to MBTD1, a subunit of the NuA4/TIP60 histone acetyltransferase (HAT) complex. In contrast to the NuA4/TIP60 complex, ZMYND11 is linked to repression of actively transcribed genes through recognition of H3.3K36me3. To understand the abnormal molecular events that expression of this ZMYND11-MBTD1 fusion protein can create, we performed its biochemical and functional characterization in comparison to each individual fusion partner. ZMYND11-MBTD1 is stably incorporated into the endogenous NuA4/TIP60 complex but does not bring any additional interactors as the fusion lacks the MYND domain of ZMYND11. Nevertheless, this truncated ZMYND11 moiety in the fusion leads to mislocalization of the NuA4/TIP60 complex on the body of genes normally bound by ZMYND11 in the genome, in a PWWP-H3.3K36me3 interaction-dependent manner. This can be correlated to increased chromatin acetylation and altered gene transcription, most notably on the *MYC* oncogene, and alternative splicing. Importantly, expression of ZMYND11-MBTD1, but not the individual fusion partners, during embryonic stem cell differentiation, leads to decreased expression of specific differentiation markers, while favoring Myc-driven pluripotency. It also favors self-renewal of hematopoietic stem/progenitor cells. Altogether, these results indicate that the ZMYND11-MBTD1 fusion protein functions primarily by mistargeting the NuA4/TIP60 complex to the body of genes, altering normal transcription of specific genes, likely driving oncogenesis in part through the Myc regulatory network.

**Highlights:**

-A recurrent chromosomal translocation detected in cannibalistic acute myeloid leukemia leads to the production of a ZMYND11-MBTD1 fusion protein.
-The ZMYND11-MBTD1 fusion protein is stably incorporated into the endogenous NuA4/TIP60 complex.
-ZMYND11-MBTD1 leads to mistargeting of NuA4/TIP60 activity to the coding region of ZMYND11-target genes, altering gene expression and splicing.
-ZMYND11-MBTD1 binds the *MYC* gene leading to its upregulation, favoring growth and pluripotency while inhibiting differentiation markers.

## INTRODUCTION

Acute myeloid leukemia (AML) is a type of hematological cancer characterized by an expansion of immature cells called blasts for undifferentiated myeloid precursors. This clonal expansion leads to impaired hematopoiesis and bone marrow failure. Although intensive chemotherapy induces a remission for many patients, relapse still occurs in some patients and the survival rate after relapse is poor (Döhner et al., 2015; Padella et al., 2019). Thus, understanding the molecular mechanism driving AML is necessary to improve treatment by identifying potential therapeutic targets that are more effective. It is already known that this cancer is due to the presence of genetic abnormalities such as somatic mutations and a significant number of gene fusions induced by chromosomal translocations (2013; Zjablovskaja and Florian, 2019). A specific recurrent translocation, t(10;17) (p15;q21), has been described over the past few years and has been linked to a cannibalistic form of AML (Plesa and Sujobert, 2019). Based on FISH analyses, this translocation creates a merge of *ZMYND11* and *MBTD1* genes (De Braekeleer et al., 2013). The break points are in exon 11 or 12 of *ZMYND11* and exon 3 of *MBTD1*, creating an in-frame coding sequence that encompasses a truncated ZMYND11 fused to a full-length MBTD1 (De Braekeleer et al., 2013; de Rooij et al., 2016; Plesa and Sujobert, 2019; Tempescul et al., 2007; Yamamoto et al., 2018) (**Fig. 1A**).

**Figure 1:**
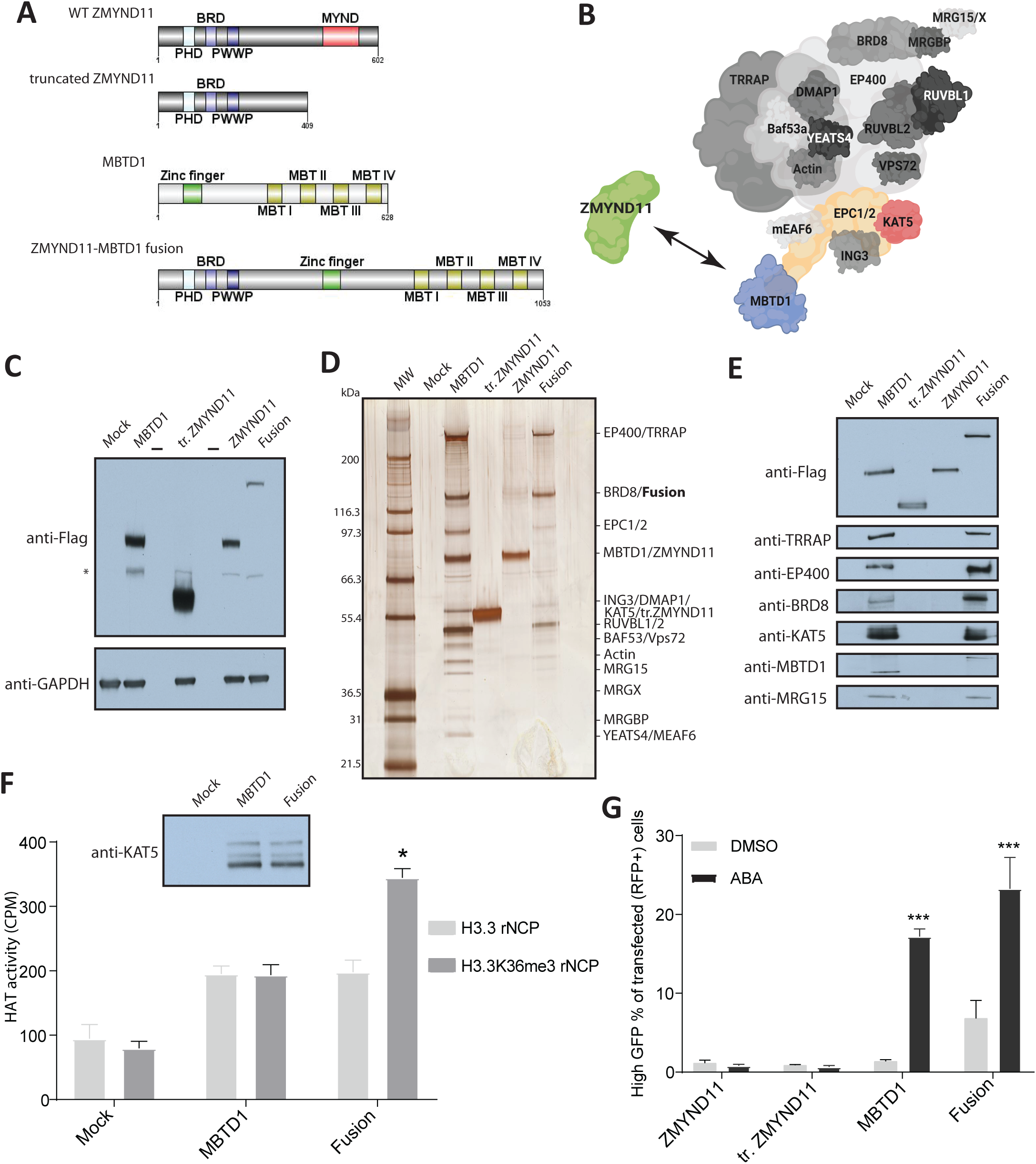
The ZMYND11-MBTD1 fusion protein is stably incorporated within the endogenous NuA4/TIP60 acetyltransferase complex. **A.** Schematic representation of MBTD1, WT ZMYND11, truncated ZMYND11 and ZMYND11-MBTD1 fusion. The loss of the C-terminal MYND domain in ZMYND11 is seen in both truncated ZMYND11 and the ZMYND11-MBTD1 fusion. **B.** Schematic representation of the NuA4/TIP60 complex showing where the physical link between MBTD1 and ZMYND11 is created by the fusion (made with BioRender). **C.** Whole-cell extract of isogenic K562 cells stably expressing MBTD1-3xFlag 2xStrep, WT ZMYND11-3xFlag2xStrep, truncated (tr)ZMYND11-3xFlag2xStrep and the ZMYND11-MBTD1 fusion-3xFlag2xStrep from the *AAVS1* safe harbor locus. Mock is a control cell line expressing an empty 3xFlag2xStrep tag (* non-specific band). **D.** Tandem affinity purifications from nuclear extracts of the cell lines shown in (C). The biotin elutions from the Strep-Tactin beads were analyzed on 4-12% SDS-PAGE followed by silver staining. Zmynd11 and subunits of the NuA4/TIP60 complex are labeled and were all identified by mass spectrometry (Table S1). **E.** Purified complexes from (D) were analyzed by western blot with the indicated antibodies to confirm the presence of known subunits of the NuA4/TIP60 complex. **F.** ZMYND11-MBTD1 incorporation into NuA4/TIP60 provides a gain of function towards H3.3K36me3. *In vitro* histone acetylation assays with purified native complexes from 1D and recombinant nucleosome core particles (rNCP). Fusion-containing NuA4/TIP60 shows an increase of HAT activity on nucleosomes methylated on H3.3K36 compared to unmodified, which is not the case for normal NuA4/TIP60 (MBTD1) (mean ± s.e.m, n=4). The embed western blot panel shows the relative KAT5 signal as control. **G.** The ZMYND11-MBTD1 fusion is a transcriptional activator similar to MBTD1, whereas ZMYND11 and its truncation are not. Transcription activation reporter assay quantified by flow cytometry analysis (Alerasool et al., 2022). It shows the % of cells transfected with ZMYND11, truncated ZMYND11 (tr.Z11), MBTD1, or ZMYND11-MBTD1 fusion that express high level of GFP upon ABA-mediated targeting of the proteins to the reporter. At least 25,000 cells were analyzed for each replicate (mean ± s.e.m, n=3). DMSO is used as a negative control. Statistical analyses were performed by two-way ANOVA test followed by Tukey’s test, *, p < 0.05.

The nuclear factor ZMYND11 (also called BS69) contains a tandem reader module of the histone variant H3.3 trimethylation on lysine 36 (H3.3K36me3), including a PWWP domain, a plant homeodomain (PHD) and a bromodomain. It also contains a C-terminal MYND domain that is important for binding proteins such as transcription factors, the spliceosome machinery, and chromatin factors (Guo et al., 2014; Sugden et al., 2019; Velasco et al., 2006; Wang et al., 2014; Wen et al., 2014). Interestingly, these interactions confer to ZMYND11 a function in pre-mRNA processing, mostly by promoting intron-retention (Guo et al., 2014). In addition, ZMYND11 acts as an unconventional transcription co-repressor by modulating RNA polymerase II elongation on highly expressed genes (Wen et al., 2014). The *ZMYND11* gene is often deleted or downregulated in many types of cancer, including breast cancer (Chen et al., 2019). Moreover, overexpression of ZMYND11 inhibits tumor cell growth, highlighting a tumor suppressor function (Chen et al., 2019; Wen et al., 2014). Recently, it was shown that, in immune cells, stimulation-induced H3.3S31 phosphorylation leads to ejection of ZMYND11, allowing rapid transcriptional activation (Armache et al., 2020). Similarly, the H3.3G34R histone mutation linked to pediatric high-grade gliomas is also linked to displacement of ZMYND11 from chromatin (Bressan et al., 2021).

MBTD1 (Malignant Brain Tumor Domain-containing protein 1) was originally identified in the Polycomb Group family of proteins because of its 4 MBT domains. Knock-out of the *MBTD1* gene in mice leads to death soon after birth caused by defects in hematopoietic stem cells and skeletal formation (Hiroaki Honda et al., 2011). MBTD1 is a stable stoichiometric subunit of the large NuA4/TIP60 protein complex (Jacquet et al., 2016; Zhang et al., 2020). NuA4/TIP60 is highly conserved in eukaryotes, harboring histone acetyltransferase activity towards H4/H2A via the haplo-insufficient tumor suppressor KAT5/Tip60 subunit as well as ATP-dependent histone exchange activity via the EP400 subunit (Pradhan et al., 2016; Steunou A-L et al., 2014). NuA4/TIP60 complex is an essential regulator of gene expression and genome stability, implicated in many cellular functions such as apoptosis and stem cell maintenance/renewal. The function of MBTD1 within NuA4/TIP60 has been linked to both gene-specific regulation and repair of DNA double-strand breaks through its MBT domains binding preference for the H4K20me1/2 histone mark (Jacquet et al., 2016; Jitka Eryilmaz et al., 2009; Zhang et al., 2020).

Well-characterized examples of fusions are found in AML, most notably translocations involving mixed-lineage leukemia 1 (MLL1) protein (KMT2A) which are found in up to 10% of acute leukemia (Winters and Bernt, 2017). These translocations have been the focus of intense research to understand their oncogenic mechanism in order to improve therapy (Birch and Shilatifard, 2020; Zhao et al., 2019). The same interest and goals logically apply to the newly described *ZMYND11-MBTD1* translocation recurrently found in AML patients. Thus, we undertook biochemical and functional analyses of the fusion protein produced by this translocation. In this report, we describe the stable integration of the ZMYND11-MBTD1 fusion protein in the full NuA4/TIP60 complex. This leads to mislocalization of NuA4/TIP60 on the body of genes normally bound by ZMYND11. Associated changes in chromatin marks, gene expression and mRNA processing are analyzed, and events likely linked to the oncogenic mechanism of the ZMYND11-MBTD1 fusion are identified.

## RESULTS AND DISCUSSION

### The ZMYND11-MBTD1 fusion protein is stably incorporated into the endogenous NuA4/TIP60 acetyltransferase complex

In order to characterize the interactome of the ZMYND11-MBTD1 fusion protein, we used genome editing to integrate a single copy of tagged cDNAs at the *AAVS1* genomic safe harbor locus (Dalvai et al., 2015). We established isogenic K562 cell lines expressing 3xFlag-2xStrep-tagged ZMYD11-MBTD1, fusion partners truncated ZMYND11(aa1-409) and full-length MBTD1, as well as full-length ZMYND11 and empty tag as controls (**Fig. 1A-B**). Expression of the different constructs from isolated cellular clones is shown in **Fig. 1C**. Tandem affinity purifications were performed from nuclear extracts using anti-Flag and Strep-Tactin beads and native elution with Flag peptides and biotin, respectively (Doyon and Côté, 2016). As we reported previously, through its stable association with EPC1, MBTD1 was purified along with all known subunits of NuA4/TIP60, as seen on silver-stained gel, western blot analysis and mass spectrometry (**Fig. 1B, D-E**, **Table S1**)(Jacquet et al., 2016; Zhang et al., 2020). ZMYND11 on the other hand was found mostly by itself but several associated substoichiometric proteins were detected, including previously linked EFTUD2 and other mRNA processing factors (**Fig. 1D-E**, **Table S1**)(Guo et al., 2014). In contrast, the truncated ZMYND11 found in the fusion lost almost all of these interactions, at least in part due to its missing MYND domain (**Fig. 1D-E**, **Table S1**)(Guo et al., 2014). Finally, purification of the ZMYND11-MBTD1 fusion protein produced an exact merge of the full NuA4-TIP60 complex with the simple addition of the truncated ZMYND11 moiety of the fusion (**Fig. 1D-E**, **Table S1**).

Since the ZMYND11 portion of the fusion protein retains its chromatin-binding domains recognizing H3.3K36me3, we tested if this leads to a gain of function for the NuA4/TIP60 complex. Indeed, in histone acetyltransferase assays with recombinant nucleosomes, the complex carrying the ZMYND11-MBTD1 fusion showed stronger acetylation of the ones carrying the H3.3K36me3 mark compared to unmodified ones, while the normal NuA4/TIP60 complex showed similar activity towards both (**Fig. 1F**). In addition, since ZMYND11 is described as a repressor of transcription, we used a recently described chemically inducible dCas9-based transcription reporter assay (Alerasool et al., 2022). In this system, clear transcriptional activation was detected in cells expressing MBTD1, as expected, but also in cells expressing the ZMYND11-MBTD1 fusion, while this was not the case with full length or truncated ZMYND11 (**Fig. 1G**). This demonstrates that the ZMYND11 moeity added to the NuA4/TIP60 complex does not impede its transcription co-activator function. Altogether, these results indicate that the ZMYND11-MBTD1 fusion is stably incorporated in the endogenous NuA4/TIP60 complex, while not bringing any significant additional interactors but providing the ability to bind the H3.3K36me3 histone mark. This suggests that the oncogenic mechanism of the fusion protein engages NuA4/TIP60 activities.

### The ZMYND11-MBTD1 fusion protein hijacks the NuA4/TIP60 complex to the coding regions of genes

Since truncated ZMYND11 moiety of the fusion protein contains strong chromatin reader motifs (PHD-Bromo-PWWP) specific for the H3.3K36me3 mark (Guo et al., 2014; Wen et al., 2014), a logical hypothesis is that it will affect NuA4/TIP60 localization in the genome. While ZMYND11 follows H3.3K36me3 on the coding region of active genes (Guo et al., 2014; Wen et al., 2014), NuA4/TIP60 is normally localized around the transcription start sites and at a subset of enhancers (Jacquet et al., 2016; Ravens et al., 2015). Thus, to elucidate the genome-wide location of the ZMYND11-MBTD1 fusion protein, we performed chromatin immunoprecipitations coupled to high throughput sequencing (ChIP-seq) using the tagged isogenic cell lines. As expected, MBTD1 and ZMYND11 bound regions showed strong enrichment for promoters and gene bodies, respectively (**Fig. 2A**). Interestingly, the truncated ZMYND11 lacking the MYND domain remained bound to almost all the same genome region as the full-length protein (**Fig. 2A-B**). This clearly indicates that ZMYND11 chromatin reader module (PHD-Bromo-PWWP) is the main if not sole determinant of ZMYND11 location in the genome. Finally, analysis of the ZMYND11-MBTD1 binding profiles showed enrichment at both promoters and gene bodies (**Fig. 2A**). But analysis of the total bound regions showed that almost all of them are shared with the truncated ZMYND11 locations (**Fig. 2B**). As expected, fusion-bound genomic locations are enriched for histone marks and transcription factors detected on active genes (**Table S2A**) while Gene ontology term (GO-term) enrichment analyses identify different biological processes, molecular functions, some pathways potentially involved in tumorigenesis (**Fig. S1A**). Interestingly, the ZMYND11-MBTD1 fusion tended to be enriched at the acceptor and donor splice sites on the coding region of genes, similar to WT ZMYND11 and truncated ZMYND11 but not MBTD1 (**Fig. S1B**). Altogether, these results indicate that the ZMYND11 moiety of the fusion drives its location in the genome, bringing along NuA4/TIP60. ChIP-seq binding profiles of ZMYND11-MBTD1 on specific genes clearly show correlation with wild type and truncated ZMYND11 on the body of these genes, supporting the concept of driving mislocalization of the NuA4/TIP60 complex along their coding region (**Fig. 2C**). Importantly, *RPSA* is naturally bound at its TSS by the NuA4/TIP60 complex, as shown here by the MBTD1 profile, while ZMYND11 and the fusion are found from the TSS to the body of the gene (**Fig. 2C**, top). Other genes of interest shown include *MYC*, *FASTK andTMUB1*, (**Fig. 2C**). This binding to the coding regions of specific genes was confirmed by ChIP-qPCR while also highlighting the very strong signals elicited by ZMYND11 and its truncated form, much less so for the fusion, most likely due to its lower expression (**Fig. 2D, 1C**)(signal at *RPSA* shown at the TSS). Since most if not all ZMYND11-MBTD1 fusion is incorporated in the NuA4/TIP60 complex (**Fig. 1D-E**), it is logical to predict that the fusion will have an impact on chromatin dynamics on the body of bound genes.

**Figure 2:**
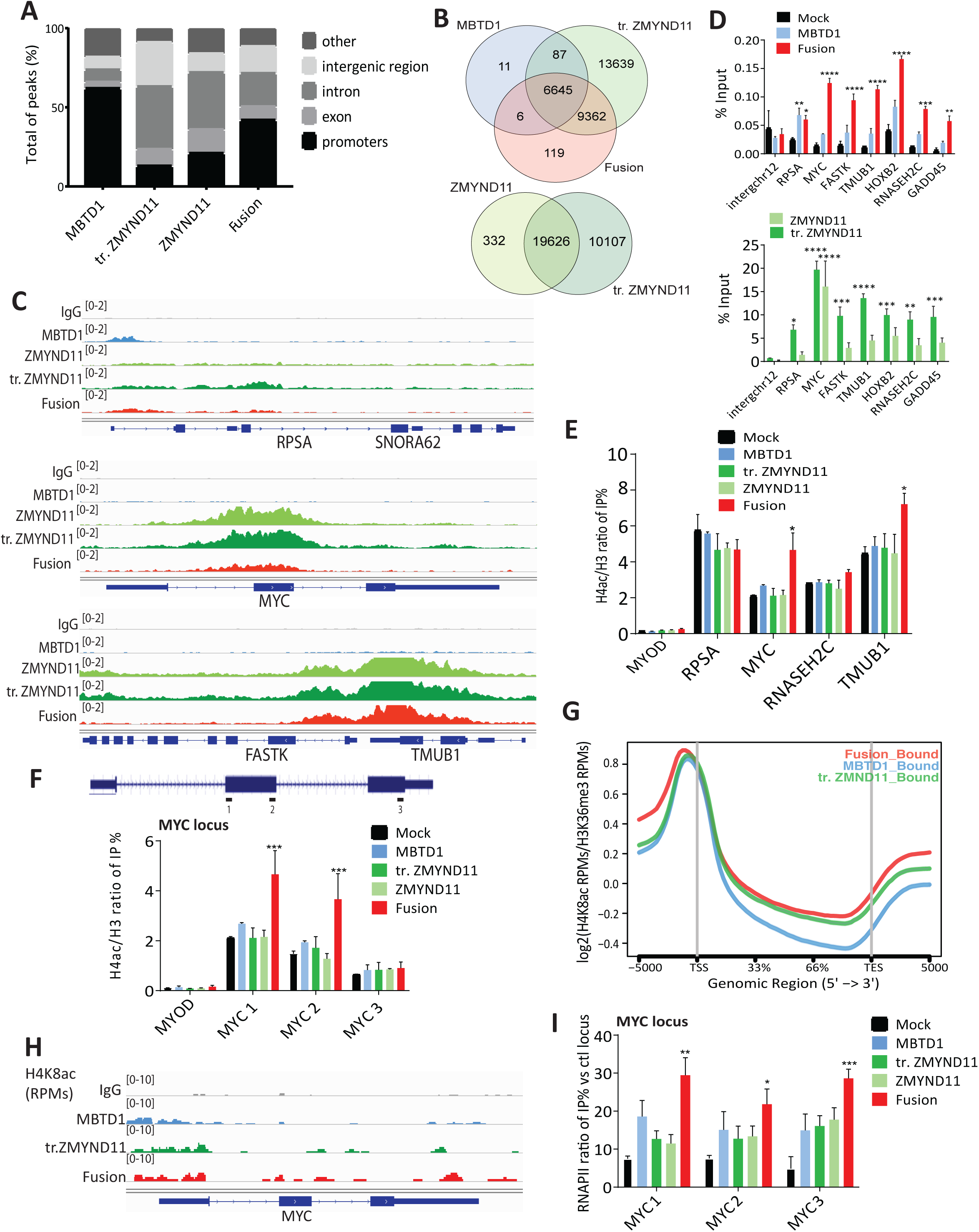
The ZMYND11-MBTD1 fusion binds ZMYND11-targets in the genome, enriched on gene bodies, affecting chromatin acetylation and RNA pol II signals on specific genes. **A.** Average of enrichment of MBTD1, WT ZMYND11, truncated (tr.) ZMYND11 and ZMYND11-MBTD1 fusion plotted to the genomic regions as promoters, exon, intron, intergenic and other (5’UTR and 3’UTR). Genome-wide localization of the fusion was analyzed by ChIP-seq and compared to data obtained in parallel with MBTD1, WT ZMYND11 and truncated (tr.) ZMYND11. Anti-Flag chromatin immunoprecipitations were performed in tagged and mock K562 cell lines used in figure 1. **B.** Comparison of genomic binding loci of truncated (tr.) ZMYND11, WT ZMYND11, MBTD1, and ZMYND11-MBTD1 fusion highlights extensive common targets between ZMYND11 and the fusion protein. Furthermore, truncated ZMYND11 can still bind the full set of ZMYND11 targets despite the loss of the MYND domain. **C.** Examples of ZMYND11-MBTD1 binding profiles on ZMYND11-target genes *RPSA*, *MYC* oncogene, *FASTK* and *TMUB1*, compared to WT and truncated ZMYND11, as well as MBTD1 and mock control. Note MBTD1 binding to the *RPSA* promoter. **D.** ChIP-qPCR confirms the localization of ZMYND11-MBTD1 on ZMYND11 target-genes (coding regions). *RPSA* promoter is used as control for the NuA4/TIP60 complex (MBTD1). Error bars represent range of two independent experiments. **E.** ChIP-qPCR of acetylated histone H4 in the different K562 cell lines. Signals are presented as a ratio of IP/input % of H4ac on total H3 to correct for nucleosome occupancy. The expression of ZMYND11-MBTD1 fusion increases the acetylation of H4 on the coding region of specific bound-genes, as *MYC* and *TMUB1*. Error bars represent the range based on two independent experiments. **F.** ChIP-qPCR of H4 acetylation along the body of the *MYC* gene, showing an increase of acetylation over the central exon specific to the cells expressing ZMYND11-MBTD1, precisely where the fusion protein is localized. The set of primers used are shown in the schematic *MYC* locus. Error bars represent the range of two independent experiments. **G.** Metagene plot showing the density of the H4K8ac/H3K36me3 ratio measured by CUT&RUN-seq across genes bound by the ZMYND11-MBTD1 fusion, MBTD1 and truncated (tr.) ZMYND11, in K562 cells (TSS: transcription start site; TES: transcription end site; RPMs: reads per million). **H.** Profile of H4K8ac sequencing reads (per million reads, RPMs) obtained by CUT&RUN over the *MYC* gene. Cells expressing the ZMYND11-MBTD1 fusion exhibit an increase of H4K8ac reads on the gene body compared to cells expressing MBTD1 or tr.ZMYND11. An IgG control is shown for background signals. **I.** ChIP-qPCR of RNA polymerase II (RNAPII) signals are presented as a ratio of IP/input % on a negative control locus (*MYOD*). Error bars represent the range of two independent experiments. Statistical analyses were performed by two-way ANOVA test followed by Tukey’s test, *, p < 0.05 , **, p < 0.01, ***, p < 0.001, ****, p <0.0001. See also Fig. S1.

### Aberrant H4ac and RNAPII enrichments on a subset of genes bound by the ZMYND11-MBTD1 fusion

To confirm that ZMYND11-MBTD1 leads to mislocalization of NuA4/TIP60 and its activity, we measured H4 acetylation by ChIP-qPCR on selected genes. While several genes did not show any significant changes of H4ac in cells expressing the fusion or its partners independently, we detected a clear increase of histone acetylation on the coding region of the *MYC* oncogene (and the *TMUB1* gene), specifically in the presence of the fusion (**Fig. 2E**). Since ZMYND11 binds to actively transcribed genes carrying the H3.3K36me3 mark, it is possible the fusion bringing NuA4/TIP60 activities has little visible impact on an already open highly dynamic chromatin, on which ZMYND11 may have little effect to start with. Nevertheless, we then focused on the clearly impacted *MYC* locus and analyzed H4ac at different locations along the gene (**Fig. 2F**). The increase of acetylation specific to the fusion-expressing cells is localized over the central exon of the MYC gene, exactly where ZMYND11 and the fusion are enriched on their ChIP-seq profiles (**Fig. 2F versus 2C**). These results clearly demonstrate that the ZMYND11-MBTD1 fusion does bring the NuA4/TIP60 complex and mis-targets its acetyltransferase activity on the coding region of the *MYC* oncogene.

Normally, histone acetylation is enriched over the promoter/TSS region where the H3K4me3 mark is located but decreases along the coding region with the appearance of the H3K36me3 mark as chromatin needs more stability to avoid spurious transcription (Barth and Imhof, 2010). We then asked if the change of H4 acetylation induced by the ZMYND11-MBTD1 fusion has an impact on these methylation marks. We could not see any fusion-specific changes on *MYC* or *RPSA*, only maybe some effect of the fusion partners by themselves (**Fig. S1C-D**). We then took the advantage of the CUT&RUN (Cleavage Under Targets & Release Using Nuclease) method to provide genome-wide view of both H3K36me3 and H4K8ac. As expected, MBTD1, truncated ZMYND11 and fusion-bound genes, in their respective expressing cell lines, showed high level of H3K36me3 signals since they are actively transcribed (**Fig. S1E**). Interestingly, genes bound by the ZMYND11-MBTD1 fusion may show on average slightly higher level of H3K36me3, but this was not obvious when looking at the specific profile on the *MYC* gene (**Fig. S1F**). In parallel, we could detect in average a slight increase of H4K8ac signals on genes bound by the fusion, even when corrected for the H3K36me3 signal on those genes (**Fig. 2G**). The H4K8ac CUT&RUN profile on the *MYC* gene also showed higher read counts on the coding region compared to controls, in agreement with the ChIP-qPCR results (**Fig. 2H**).

Next, we looked at the density of RNA polymerase II (RNAPII) as local changes in chromatin dynamics due to increased acetylation may affect transcription elongation. Total RNAPII measured by ChIP-qPCR along the *RPSA* gene showed a strong increase of signal specifically at the TSS in cells expressing the fusion (**Fig. S1G**). Interestingly, this was also seen in cells expressing the truncated ZMYND11 but not the full-length protein (**Fig. S1G**). Thus, these effects seem independent of NuA4/TIP60 and its HAT activity (**Fig. 2E**) and are likely produced somehow by the loss of the MYND domain in ZMYND11. In contrast, RNAPII signals were specifically increased by the ZMYND11-MBTD1 fusion at 3 locations along the *MYC* gene compared to control cell lines (**Fig. 2I**). These results suggest that mis-targeting of NuA4/TIP60 activity by the ZMYND11-MBTD1 fusion can lead to increased transcription on genes like the *MYC* oncogene and this could be part of the oncogenic mechanism.

### H3.3K36me3-binding by the ZMYND11-MBTD1 fusion is critical for mis-targeting the NuA4/TIP60 complex

The N-terminal module of ZMYND11 containing tandem PHD, Bromo and PWWP domains is essential for its binding to chromatin, recognizing the histone variant H3.3 methylated on lysine 36 (H3.3K36me3), a mark associated with the body of actively transcribed genes (Guo et al., 2014; Wen et al., 2014). As shown above, this binding can increase the HAT activity of a NuA4/TIP60 complex containing the ZMYND11-MBTD1 fusion *in vitro* (**Fig. 1F**). To determine the significance of this gain-of-function *in vivo*, we produced point mutations in the PWWP domain of ZMYND11 within the fusion. Residues W294 and F310 were mutated as they are already known to be crucial for H3.3K36me3-binding (Wen et al., 2014). Wild-type and mutant fusion proteins were then stably expressed from the AAVS1 locus in K562 cells (**Fig. 3A**). ChIP-qPCR were then performed to evaluate mutant effects on genomic localization of the fusion and its function. Both W294A and F310A single mutants drastically disrupted the binding of the fusion protein to the *MYC* oncogene and *TMUB1* (**Fig. 3B**). This was also correlated with a clear loss of H4 acetylation driven by the wild-type fusion on the body of these genes (**Fig. 3C**). These results clearly demonstrate that mis-targeting of NuA4/TIP60 by ZMYND11-MBTD1 fully depends on its recognition of H3.3K36me3 on the on the body of active genes, implicating this recognition mechanism in the leukemogenesis driven by the fusion.

**Figure 3:**
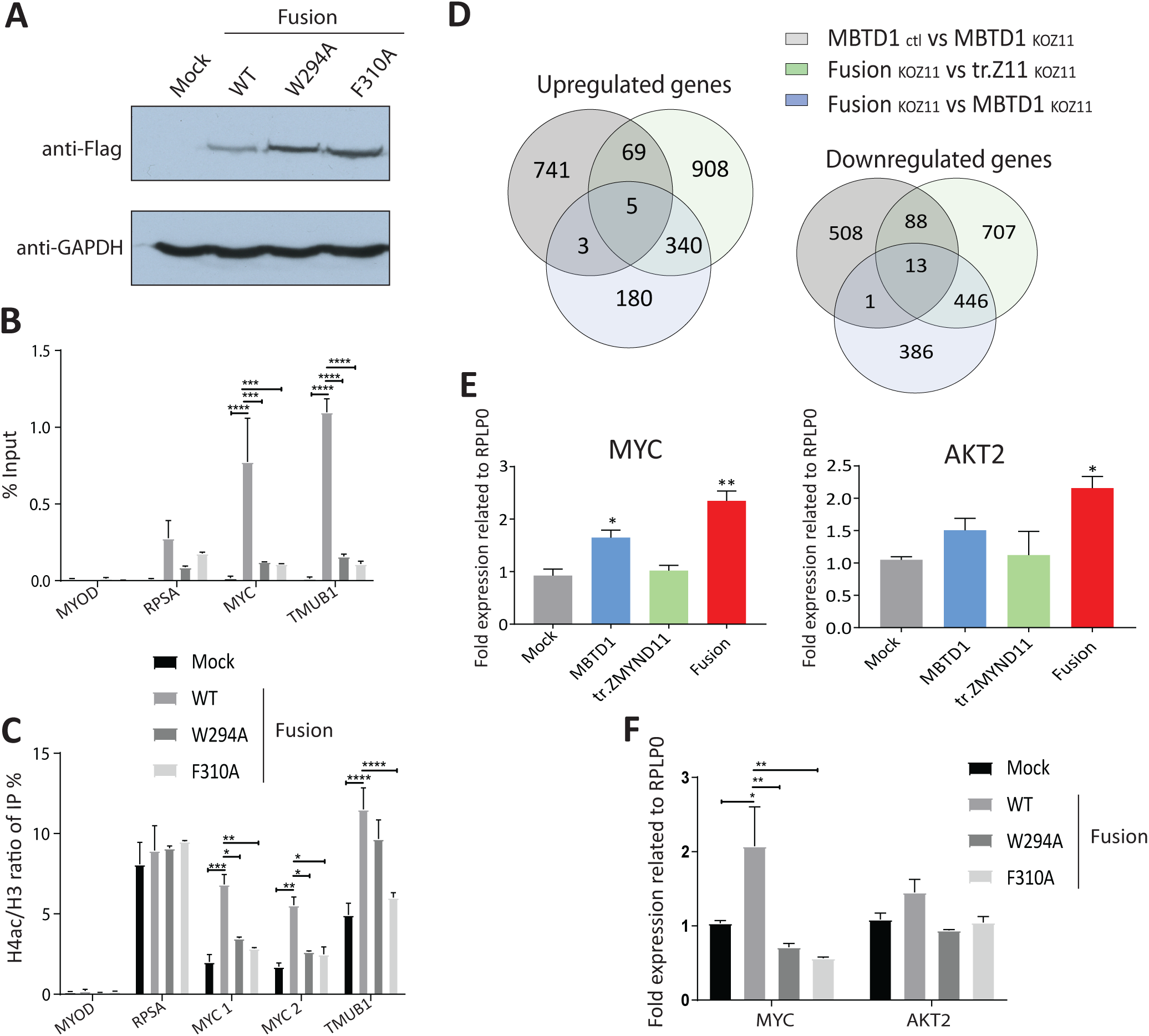
Mistargeting of NuA4/TIP60 by the ZMYND11-MBTD1 fusion and its effect on gene expression depend on the H3.3K36me3-binding PWWP domain of ZMYND11. **A.** Western blot of whole-cell extracts from isogenic K562 cells stably expressing the WT ZMYND11-MBTD1 fusion-3xFlag2xStrep or PWWP single mutants W294A or F310A from the AAVS1 safe harbor locus. A mock control cell line expressing an empty 3xFlag2xStrep tag is also shown. **B.** Anti-Flag ChIP-qPCR quantifying binding of the ZMYND11-MBTD1 fusion in K562 cells (from A) stably expressing either WT or H3.3K36me3-binding defective PWWP mutants (W294A or F310A), at the indicated genes. Mock cell line and the *MYOD* promoter are used as negative controls. **C.** ChIP-qPCR showing that the increase of H4 acetylation on the body of *MYC* and *TMUB1* genes driven by the ZMYND11-MBTD1 fusion is lost in cells expressing H3.3K36me3-binding mutants (W294A and F310A). The *RPSA* promoter shows no change of H4 acetylation levels while *MYOD* and a mock cell line are used as controls. **D.** 340 genes are found upregulated (left) and 446 downregulated (right) upon expression of the ZMYND11-MBTD1 fusion when compared to both cell lines expressing truncated ZMYND11 or MBTD1, over endogenous *ZMYND11* gene knock-out (KOZ11, by CRISPR/Cas9) as well as compared to control wild type cells (long reads RNA-seq, p-value 0.01% and 25% FDR). **E.** RT-qPCR of *MYC* and *AKT2* expression confirming that they are specifically upregulated in in K562 cells expressing the ZMYND11-MBTD1 fusion compared to partners (depleted in endogenous ZMYND11 as in D). **F.** RT-qPCR of *MYC* and *AKT2* genes in cells expressing the ZMYND11-MBTD1 fusion showing that H3.3K36me3-binding PWWP mutants (as in A) fail to show increased transcription. Error bars represent the range of two independent experiments. Statistical analyses were performed by two-way ANOVA test followed by Tukey’s test, *, p < 0.05 , **, p < 0.01, ***, p < 0.001, ****, p <0.0001.

### The ZMYND11-MBTD1 fusion protein alters the expression of several genes linked to growth and differentiation

To get a bigger picture of the effect of ZMYND11-MBTD1 on gene expression, we use CRISPR/Cas9 to knock-out the endogenous *ZMYND11* gene (Agudelo et al., 2017) in our K562 cell lines expressing the fusion protein or its partners from the *AAVS1* locus. We used two gRNAs to induce a truncation in the C-terminal MYND domain of endogenous ZMYND11 and isolated 2 independent clones (**Fig. S2A-C**). We kept the MBTD1^AAVS1^ cell line that still expressed endogenous ZMYND11 as a control. Taking advantage of the Oxford Nanopore Technologies long-read Sequencing, a nanopore-based single-molecule sequencing that is more informative for gene isoform/splice variant characterization (Jason L Weirather et al., 2017), we sought to determine changes in mRNA levels upon expression of the ZMYND11-MBTD1 fusion compared to its fusion partners. Differential mRNA expression was investigated comparing sequences between the fusion cell lines (endogenous ZMYND11 KO + ZMYND11-MBTD1^AAVS1^) and three control cell line pairs (endogenous ZMYND11-wild type + MBTD1^AAVS1^ (control), ZMYND11-KO + MBTD1^AAVS1^ (MBTD1) and ZMYND11-KO + truncated ZMYND11^AAVS1^ (tr.ZMYND11)). 340 genes were found upregulated and 446 downregulated in the cells expressing the ZMYND11-MBTD1 fusion compared to both cell lines expressing fusion partners individually (**Fig. 3D**). Interestingly, gene ontology analyses show that the genes upregulated are enriched for ribosome biogenesis, hence supporting growth/proliferation, and downregulated genes are strongly enriched for biological processes involved in myeloid cell activation, which likely reflects K562 cells undergoing some dedifferentiation programming, a hallmark of oncogenesis (**Fig. S2D**). Importantly, the correlation between ChIP-seq and RNA-seq data shows that the majority (∼2/3) of upregulated genes are bound by the fusion (**Fig. S2E**). Thereafter, we used RT-qPCR to validate 2 genes upregulated and important positive regulators of cell growth/proliferation, *MYC* and *AKT2*, and confirmed their increased transcription in cells expressing the ZMYND11-MBTD1 fusion compared to the control cell lines (**Fig. 3E**). Interestingly, a significant portion of the genes affected by the fusion are known to be regulated by MYC/MAX transcription factors, suggesting indirect effect on their transcription due to increased expression of MYC (**Table S2B**). Importantly, using the PWWP/H3.3K36me3 - binding ZMYND11-MBTD1 mutants, we observed the loss of increased *MYC* and *AKT2* expression driven by the wild type ZMYND11-MBTD1 fusion (**Fig. 3F**). Altogether, these results argue that expression of the ZMYND11-MBTD1 fusion protein and its association to NuA4/TIP60 lead to transcription changes of several H3.3K36me3-marked genes normally bound by ZMYND11, favoring cell proliferation/growth and inhibiting differentiated phenotypes.

### The ZMYND11-MBTD1 fusion protein disrupts gene-specific ratio of mRNA splice variants

Almost 95% of human mRNAs are alternatively spliced and many studies already proved that chromatin marks affect splicing (for review (Hnilicova and Stanek, 2011)). Moreover, ZMYND11 itself was shown to interact with splicing factors and directly affect splicing events, especially intron retention (Guo et al., 2014). However, we have shown above that ZMYND11 normal interactome was lost in the fusion and that local chromatin structure can be affected by mislocalization of NuA4/TIP60 on ZMYND11-targeted gene bodies (**Fig. 1, 2**). It was then important to determine whether the fusion protein perturbs normal mRNA splicing events causing different gene isoforms to be expressed. Therefore, we sought to correlate changes in isoform expression to their respective gene changes in the context of overexpression of the fusion or control partners. As expected, we found that many of the differentially expressed transcripts (9040) had a similar expression pattern as their respective genes (**Fig. S2F**, left heatmap). However, 214 transcripts that were differentially expressed at both the gene and transcript level instead showed transcript expression that did not match their respective genes (**Fig. S2F**, right heatmap). We then focused on transcripts that are not differentially changed at the gene level (**Fig. S2F** left). Interestingly, we also identified a total of 5634 transcripts (∼38% of differentially expressed transcripts) where differential expression was detectable at the isoform level but not at the gene level (**Fig. 4A**). This suggests that, for an important number of genes, alternative isoform choice is occurring between samples. Analysis of these differentially expressed transcripts (a total of 5634 + 214) highlighted 111 high-confidence transcripts that were affected by overexpression of the fusion protein when comparing to overexpression of either control partners (**Fig. 4B**). Furthermore, genome-wide analysis of splicing events revealed that the fusion led to a significant decrease in exon skipping events and an increase in intron retention events compared to the MBTD1 control partner (**Fig. 4C**). In contrast, no significant change in splicing events was observed when the fusion was compared to truncated ZMYND11 control (**Fig. 4C**). Importantly, almost 80% of the corresponding genes are bound by the ZMYND11-MBTD1 fusion and 64% are already known to be alternatively spliced (**Table S2C**). This was further confirmed by RT-qPCR of two selected genes in which the cells expressing the fusion protein specifically showed a change in the ratio of splice variants compared to its partners and control cell lines (**Fig. 4D-E**). Altogether, these data demonstrate prominent specific effects of the ZMYND11-MBTD1 fusion on gene expression and transcript isoforms levels, part of the mechanism that is likely at play in acute myeloid leukemia harboring this chromosomal translocation.

**Figure 4.**
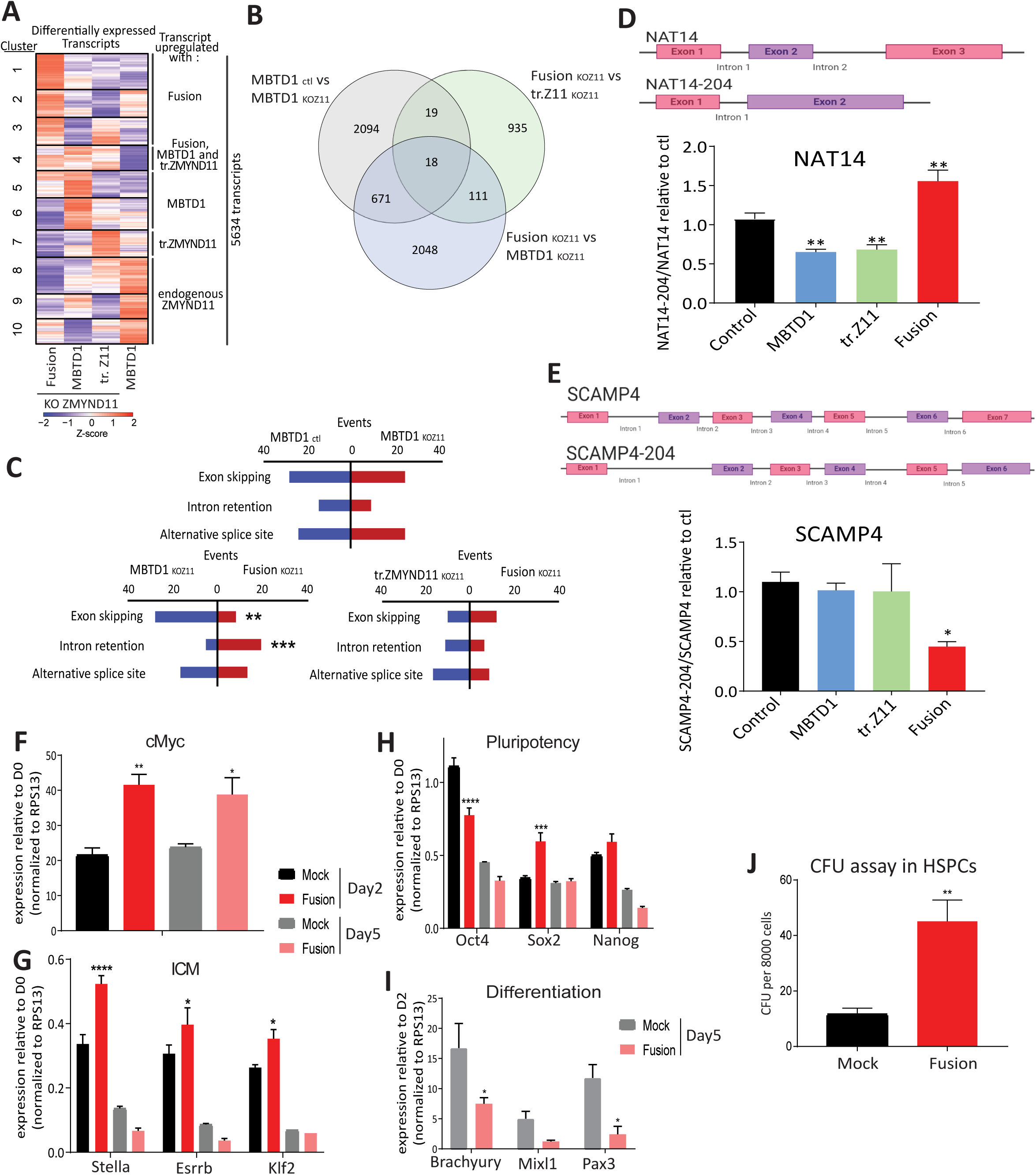
Expression of the ZMYND11-MBTD1 fusion alters splicing events and favors Myc-driven pluripotency. **A.** Heatmap of expression (in Z-scores of Log_2_(CPM)) for transcripts where differential expression was detectable at the isoform level but not at the gene level, upon expression of the ZMYND11-MBTD1 fusion protein, tr. (truncated) ZMYND11 or MBTD1 in KO ZMYND11 (KOZ11) K562 cell lines, or between MBTD1 in KO ZMYND11 cells and control MBTD1 cells with wild-type endogenous ZMYND11. Transcripts are hierarchically clustered, and clusters of similarly expressed transcripts in specific samples are indicated. **B.** Venn diagram showing overlaps in differentially expressed transcripts driven by the fusion compared to control cell lines. **C.** Transcriptome-wide analysis of the occurrence of splicing events between long-read RNA-seq samples. Exon skipping, intron retention and alternative splice site choice events are shown. Significant divergence from expected splicing event counts was assessed by Chi-square test of independence (**: p-value ≤ 0.01, ***: p-value ≤ 0.001). **D-E**. Canonical and spliced isoforms of NAT14 and SCAMP4 mRNAs measured by RT-qPCR and normalised to RPLP0. NAT14-204 transcript and SCAMP4-204 transcript are upregulated and downregulated, respectively, in cells expressing the fusion compared to control cell lines. Values represent means relative to control cell lines from two biological replicates and error bars are range. Statistical analyses were performed by two-way ANOVA test followed by Tukey’s test, *, p < 0.05, **, p < 0.01, ***, p < 0.001. **F-I**. Relative transcriptional levels of indicated genes upon inducible expression of the ZMYND11-MBTD1 fusion in mouse embryonic stem cells (mESCs) that were induced for differentiation for 5 days. Mock is used as control. Markers of both pluripotency and differentiation are indicated and normalized to D0 or D2, respectively. The values are presented as mean ± SD of three independent experiments. Statistical analyses were performed by two-tailed t-test for c-Myc and two-way ANOVA test followed by Tukey’s test, *, p < 0.05, **, p < 0.01, ***, p < 0.001, for other markers. **J.** Quantification of colony-forming units (CFU) assay performed with hematopoietic stem/progenitor cells (HSPCs) transduced with empty vector (Mock) or ZMYND11-MBTD1 fusion (mean ± s.e.m, 9 replicates across 3 biological replicates). T-test was performed with a p-value=0.004.

### The ZMYND11-MBTD1 fusion protein favors growth and inhibits expression of differentiation markers

ZMYND11 and KAT5/Tip60 are already known to have a tumor suppressor function and to be downregulated in many human cancers (Gorrini et al., 2007; Wen et al., 2014). We first expressed the fusion, its partners and empty vector in mouse embryonic stem cells (mESCs) (**Fig. S2G**) undergoing spontaneous differentiation. RT-qPCR was used to monitor the expression of different sets of genes linked to pluripotency and early differentiation markers. Expression of *Pax3* (neural epithelium and early neural crest) was downregulated by the over-expression of ZMYND11-MBTD1 fusion, while pluripotency markers *Stella* (*Dppa3*/*Pgc7*) and *Myc* were upregulated (**Fig.S2H**). These results suggest that expression of the fusion protein inhibited differentiation of ESCs, which is consistent with its oncogenic properties. We then generated stable mESCs with inducible expression of the fusion construct and induced differentiation for 5 days. An increase of *c-Myc* expression in the presence of ZMYND11-MBTD1 was detected after 2 and 5 days (**Fig. 4F**). Other pluripotency markers also increased after 2 days, such as Inner Cell Mass (ICM) markers, including *Stella*, *Sox2, Esrrb,* and *Klf2* (**Fig. 4G-H**). In contrast, pluripotency markers *Nanog* and *Oct4* were not affected or decreased, respectively (**Fig. 4H**). Moreover, differentiation markers *Brachyury* and *Pax3* were significantly decreased after 5 days in the presence of the fusion (**Fig. 4I**). Importantly, Stella, a H3K9me reader and inhibitor of DNA demethylation, has been implicated in a Myc-driven self-reinforcing regulatory network that maintains mESC identity, independently of Sox2 and Oct4 (Fagnocchi et al., 2016), indicating that ZMYND11-MBTD1 binding to the coding region of the *MYC* gene and increasing its expression may be a major oncogenic mechanism for this specific leukemia. Finally, to determine if the ZMYND11-MBTD1 fusion can act as a driver of leukemic features, we assessed its transforming potential in healthy hematopoietic stem/progenitor cells (HSPCs) isolated from mouse bone marrow. Stable ectopic expression of ZYND11-MBTD1 robustly promoted the production of primitive progenitors compared to empty vector transduced (Mock) controls as measured by an increase of colony forming units (**Fig. 4J**). These data clearly support ZMYND11-MBTD1 as a driver of oncogenic transformation, negating the natural tumor suppressor functions of ZMYND11 and NuA4/TIP60.

## Conclusions

The recurrent chromosomal translocation t(10;17) (p15;q21) found in cannibalistic acute myeloid leukemia leads to the production of a fusion protein linking two factors with tumor suppressor activity, ZMYND11 and MBTD1. This physical merge disrupts ZMYND11 interactome with the mRNA processing machinery but not its strong epigenetic reader affinity for H3.3K36me3 located on the coding region of transcribed genes. As the fusion protein is stably integrated into the NuA4/TIP60 complex, it leads to mislocalization of this important chromatin modifier/remodeler complex to the body of genes, altering chromatin dynamics, transcription elongation and co-transcriptional mRNA processing events on specific genes (**graphical abstract**). In contrast to its fusion partners, ZMYND11-MBTD1 favors cell growth/proliferation, likely in part through expression of the *MYC* oncogene, and inhibits expression of specific differentiation markers. While this work was processed for submission, highly related findings were reported, using expression of the fusion protein in murine HSPCs to show the importance of Tip60 interaction and downstream histone acetylation for leukemogenesis (Li et al., 2021). Furthermore, the onco-histone mutation H3.3G34R in pediatric high grade glioblastoma has been linked to impaired recruitment of ZMYND11 and its repressor function on highly expressed genes, promoting tumorigenesis by stabilizing expression of key progenitor genes (Bressan et al., 2021), which is a mechanism reminiscent of the one described for the fusion. As mistargeting of the NuA4/TIP60 complex seems to be the main mechanism employed by the ZMYND11-MBTD1 fusion, it can be most interesting to test some specific KAT5/Tip60 HAT inhibitors (Gao et al., 2014) (Coffey et al., 2012) to determine if de novo acetylation at coding regions is indeed the main oncogenic driving force in this specific group of acute myeloid leukemia, as suggested by the parallel study (Li et al., 2021). But for potential therapeutic approaches, inhibiting the HAT activity of KAT5/Tip60 may have large pleiotropic effects due to the essential roles of NuA4/TIP60 in several essential nuclear processes. Since we recently described the structural determinants by which MBTD1 associates with NuA4/TIP60 through the EPC1 subunit (Zhang et al., 2020), an alternative more specific approach would be to design small molecules that disrupt this interaction and, in the present case, detach the ZMYND11-MBTD1 fusion protein from the NuA4/TIP60 complex in these leukemic cells.

## Acknowledgements

We thank members of the Côté laboratory for helpful discussion during the course of this project, and the Proteomics Center at CHU de Quebec Research Center for protein identification by mass spectrometry. This work was supported by grants from the Canadian Institutes of Health Research (CIHR) to J.C. (FDN-143314) and S.M.I.H. (PJT-378019), and from the Natural Sciences and Engineering Research council of Canada (NSERC) to A.F.-T. (RGPIN-2016-05844). V.F. is a recipient of a training award of the Fonds de Recherche du Québec-Santé (FRQ-S). G.K. is a recipient of a Pierre J. Durand scholarship from the faculty of Medicine of Laval University and Luc Bélanger prestige scholarship of the CHU de Québec Foundation. A.F.-T. is a tier 2 Canada Research Chair in Molecular Virology and Genomic Instability and is supported by the Foundation J.-Louis Lévesque. S.M.I.H. is a Junior 1 Research Scholar of the Fonds de Recherche du Québec-Santé. J.C. holds the Canada Research Chair in Chromatin Biology and Molecular Epigenetics.

## AUTHOR CONTRIBUTIONS

M.D., S.M.I.H. and J.C. designed the research. M.D., V.F., G.K., J.X., N.A., M.G., N.W., G.B. performed experiments. G.K. performed bioinformatic analyses. A.F.-T., M.T., K.H., S.M.I.H. and J.C. supervised work and secured funding. M.D. and J.C. wrote the manuscript with help from co-authors.

## DECLARATION OF INTERESTS

The authors declare no competing interests.

## STAR METHODS

### LEAD CONTACT AND MATERIALS AVAILABILITY

Further contact information and requests for resources and reagents should be directed to and will be fulfilled by the Lead Contact, Jacques Cote.

This study did not generate new unique reagents.

#### Data and code availability

All MS files generated as part of this study were deposited at MassIVE (http://massive.ucsd.edu). The MassIVE ID is MSV000087226 and the MassIVE link for download is https://massive.ucsd.edu/ProteoSAFe/dataset.jsp?task=50f6c31003a5463eac7919c03c5713dd.

The password for download prior to final acceptance is “ZMYND11-MBTD1”.

Raw sequences of ChIP-sequencing and long reads RNA-sequencing in K562 cells were deposited in the GEO database under accession number **GSE171673**.

### EXPERIMENTAL MODEL AND SUBJECT DETAILS

The plasmid DNAs were amplified in *E. coli* DH10b cells, in LB medium.

Cell culture: K562 and U2OS cells were obtained from the ATCC and maintained at 37 °C under 5% CO2. K562 were cultured in RPMI medium supplemented with 10% newborn calf serum and 1% GlutaMAX. U2OS cells were cultured in DMEM medium supplemented with 10% fetal bovine serum (FBS).

R1 mouse embryonic stem cells (ESCs) were a gift from Dr Andras Nagy’s laboratory and maintained at 37 °C under 5% CO2. R1 were cultured on 0.1% gelatin (Sigma-Aldrich #G1890) in 2i-LIF medium (LIF: Leukemia Inhibitory Factor at 20 ng/mL, 3µM CHIR99021 (Cedarlane Labs #S2924), and 1µM PD0325901 (Cedarlane Labs #S1036) in N2B27 media). N2B27 media consists of a 1:1 mixture of DMEM/F-12 media mix (ThermoFisher Scientific #11330057) supplemented with N2 supplement (ThermoFisher Scientific #17502048) and neurobasal media (ThermoFisher Scientific #21103049) supplemented with 1X B27 supplement (Thermo Fisher Scientific #17504044), 1 mM sodium pyruvate, 2 mM L-glutamine, and ß-mercaptoethanol (10^-4^ M).

### METHOD DETAILS

#### Production of rNCP

Bacterial expression vectors for histones H2A, H2B, H3.1 and H3.3 were purchased from Addgene #42634, #42630, #42631 and #42632, respectively. Plasmid to express X. laevis H4 in pET3a was obtained from C. Arrowsmith.

Recombinant nucleosome core particles (rNCP) have been produced from recombinant histones as previously described (Dyer, Edayathumangalam et al. 2004) (Galloy, Lachance et al. 2021). Briefly, after preparation of inclusion bodies, the histones were purified under denaturing conditions on a 5 ml HiTrap SP FF (Cytiva) cation exchange column. Fractions containing the purified histones were pooled and extensively dialysed into water and 2mM β-mercaptoethanol before lyophilization. Octamers were refolded by mixing the four histones in equimolar ratios, followed by dialysis into 2M NaCl, 10 mM Tris pH7.5, 1 mM EDTA, and then purified on a Superdex 200 HiLoad 16/600 size exclusion column (Cytiva). Recombinant nucleosome core particles (rNCPs) were reconstituted as described (Galloy, Lachance et al. 2021)(Dyer, Edayathumangalam et al. 2004). The 601 Widom DNA sequence with a 72 bp linker used to wrap the mononucleosomes was a gift from B. Bartholomew. DNA fragments were PCR amplified with the following biotinylated primers (FW: 5’-gctcggtactcgggttcaat-3’ and REV: 5’-ctccgtgcaggtcgactcta-3’), precipitated with phenol-chloroform and purified with columns (Favorgen), according to the manufacturer’s instructions. Native gel analysis was used to evaluate the quality of the reconstitution.

#### Establishment of isogenic cell lines from the *AAVS1* safe harbor

K562 cells were used to isolate clones after genome editing to stably express from the *AAVS1* safe harbor various 3xFlag2xSrep-tagged proteins from the moderately active human *PGK1* promoter, including MBTD1, ZMYND11, truncated ZMYND11 [aa1-409], and the ZMYND11-MBTD1 fusion, as described previously (Dalvai, Loehr et al. 2015).

#### Affinity purification of complexes

Native complexes were purified essentially as described before (Dalvai, Loehr et al. 2015). Briefly, Nuclear extracts were prepared from 2.5.10^9^ cells and pre-cleared with CL6B sepharose beads. FLAG immunoprecipitations with anti-Flag agarose affinity resin (Sigma M2) were performed followed by two elutions with 3xFLAG peptide in elution buffer (20 mM HEPES-KOH [pH 7.9], 10% glycerol, 150 mM KCl, 0.1% Tween 20, 1mMDTT, 1 mM PMSF, 2 mg/mL Leupeptin, 5 mg Aprotinin, 2 mg/mL Pepstatin, 10 mM Na-butyrate, 10 mM b-glycerophosphate) with 200 ug/mL 3xFLAG peptide (Sigma). Then, STREP immunoprecipitations with Strep-tactin sepharose beads (Cedarlane) were performed followed by two elutions with elution buffer supplemented by 4mM biotin. Typically, 20µL of the first elution Strep was loaded on Nu-PAGE 4%–12% Bis-Tris gels (Invitrogen) and analyzed via silver staining.

#### Mass spectrometry analysis

Affinity purified materials were resolved on a short SDS-PAGE gel, in-gel digested with trypsin, and subsequently samples were analyzed by nano LC/MSMS either with a 5600 triple TOF or an Orbitrap Fusion instrument at the proteomics platform of CHU de Québec-Université Laval Research Center. Criteria for protein identification: Scaffold (version Scaffold_4.8.7, Proteome Software Inc., Portland, OR) was used to validate MS/MS-based peptide and protein identifications. Peptide identifications were accepted if they could be established at greater than 8.0% probability to achieve an FDR less than 1.0% by the Scaffold Local FDR algorithm.

#### Antibodies

The following antibodies were used for Western blotting at the indicated dilution: anti-FLAG -HRP conjugate (Sigma M2 A8592-1MG, 1:5000); anti-MBTD1 (Abcam ab116361, 1:1000); anti-Brd8 (Bethyl A300-219A, 1:10000); anti-P400 (Abcam ab5201, 1:1000); anti-TRRAP (bethyl A301-132A, 1:2000); anti-MRG15 (Active Motif 39361, 1:1000); anti-Tip60 (Ab 137518, 1:1000); anti-GAPDH (ThermoFisher, 39-8600-ZG003, 1/25000).

#### ChIP-seq experiments

For Flag ChIP, 1mg of cross-linked chromatin from K562 cells was incubated with 10µg of anti-Flag antibody (Sigma, M2) pre-bound on 300 µl of Dynabeads Prot-G (Invitrogen) overnight at 4°C. The beads were washed extensively and eluted in 0.1% SDS, 0.1M NaHCO3. Crosslink was reversed with 0.2M NaCl and incubation overnight at 65C. Samples were treated with RNase and Proteinase K for 2h and recovered by phenol-chloroform and ethanol precipitation.

Libraries for sequencing were prepared with TruSeq LT adaptors (Illumina). Samples were sequenced by 50 bp single reads on HiSeq 4000 platform (Illumina).

#### CUT&RUN

Cleavage Under Targets & Release Using Nuclease (CUT&RUN) was performed according to the EpiCypher CUTANA CUT&RUN Protocol. In brief, 0.5 million cells were harvested, washed, and crosslinked in 0.1% formaldehyde for 1min. Then, 125mM of glycine was added to quench cross-linking. Cells were re-suspended in wash buffer (20 mM HEPES, pH 7.5, 150 mM NaCl, 0.5 mM Spermidine, 1% Triton X-100, 0.05% SDS and 1x Protease Inhibitor cocktail (sigma)), followed by incubation at RT for 10 min with activated ConA magnetic beads (Bangs Laboratories, cat# BP531) which were washed and re-suspended in bead activation buffer (20 mM HEPES, pH 7.9, 10 mM KCl, 1 mM CaCl_2_, 1 mM MnCl_2_). After binding to activated beads, cells were permeabilized in antibody buffer (wash buffer + 0.05% digitonin + 2 mM EDTA) and incubated with IgG (Epicypher 13-0042) or H4K8ac (Abcam ab45166) or H3K36me3 (Abcam ab9050) antibody (1:100 dilution) on nutator overnight at 4°C. On the next day, the cell-bead slurry was washed twice with cold digitonin buffer (wash buffer + 0.01% digitonin) and then incubated with pAG-MNase (2.5ml, EpiCypher, Cat# EP151016) for 10 min at RT, followed by two washes with cold digitonin buffer. MNase was then activated by addition of CaCl2 to cleave targeted chromatin for 2 h at 4 °C. To stop Mnase activity, chromatin was incubated 10min at 37°C in stop elution buffer (340 mM NaCl, 20 mM EDTA, 4 mM EGTA, 50 μg/mL RNase A, 50 μg/mL Glycogen). DNA fragments were then incubated overnight at 55°C after added 0.8ml of 10% SDS and 2ul of 20mg/ml proteinase K. The DNA was purified from the supernatant using the NEB Monarch DNA Cleanup Kit (NEB, cat# T1030) per manufacturer’s instruction. Finally, 5–10 ng of purified CUT&RUN-enriched DNA was used to prepare Illumina library using the NEB Ultra II DNA Library Prep Kit per manufacturer’s instructions.

#### Processing, alignment and peak calling of ChIP-seq and CUT&RUN-seq data

FastQ format reads were aligned to the hg19 human reference using the Bowtie alignment algorithm (Langmead et al., 2009). Bowtie2 version 2.1.6 was used with the pre-set sensitive parameter to align ChIP sequencing and CUT&RUN sequencing reads. MACS version 2.0.10 (model-based analysis of ChIPseq) peak finding algorithm was used to identify regions of ChIP-Seq or CUT&RUN-seq enrichment over the background (Feng et al., 2012). The pipeline, commands, and parameters that were used are: Trimming of sequence (filter out 39 adaptor and remove last 2 bases and 3 extra bases if it matches with adaptor sequence). Mapping sequences to human genome (hg19) using Bowtie: (i) command: bowtie2 -p 8 –sensitive -x genome/genome -U sequence.reads.fastq – S sample.sam. Peak calling algorithm MACS: (i) command : macs2 callpeak -t ChIPseq.data.bam -c input.sample.file.bam --broad -f BAM -g hs -n [directory] –outdir MACS2_files --nomodel --shiftsize 100 –B. Unique mapped read values were normalized to library size. Peaks were annotated as per human genome EnsDb.Hsapiens.v75 – hg19.

#### Chip qPCR

ChIP was performed according to the protocol described in (Avvakumov, Lalonde et al. 2011). Briefly, for anti-FLAG ChIP, 1 mg of cross-linked chromatin from K562 cells was incubated with 10 mg of anti-FLAG antibody (Sigma, M2) oranti-IgG antibody (Millipore-PP64) pre-bound on 300 ml of Dynabeads Prot-G (Invitrogen) overnight at 4°C. The beads were washed extensively and eluted in 0.1% SDS, 0.1 M NaHCO3. Crosslink was reversed with 0.2 M NaCl and incubation overnight at 65°C.

Samples were treated with RNase and Proteinase K for 2 h and recovered by phenol chloroform extraction and ethanol precipitation. Also, 200 μg of chromatin were immunoprecipitated using 1-3μg of the following antibodies: anti-H4penta-Acetyl (Upstate 06-946), anti-H3 (Abcam ab1791), anti-H3K4me3 (Abcam Ab8580), anti-H3K36me3 (Abcam Ab9050), anti-RNAPII (8WG16 Covance), overnight at 4C and incubated with 50ul of Dynabeads Protein A/G (Invitrogen) 4h at 4C. The beads were washed extensively and eluted in 0.1% SDS, 0.1M NaHCO3. Crosslink was reversed with 0.2M NaCl and incubation overnight at 65C. Samples were treated with RNase and Proteinase K for 2h and recovered by phenol-chloroform and ethanol precipitation. Quantitative real-time PCRs were performed on a LightCycler 480 (Roche) with SYBR Green I (Roche) to confirm the specific enrichment at defined loci compared to intergenic regions or negative locus. IgG-ChIPs were subtracted from FLAG-ChIP.

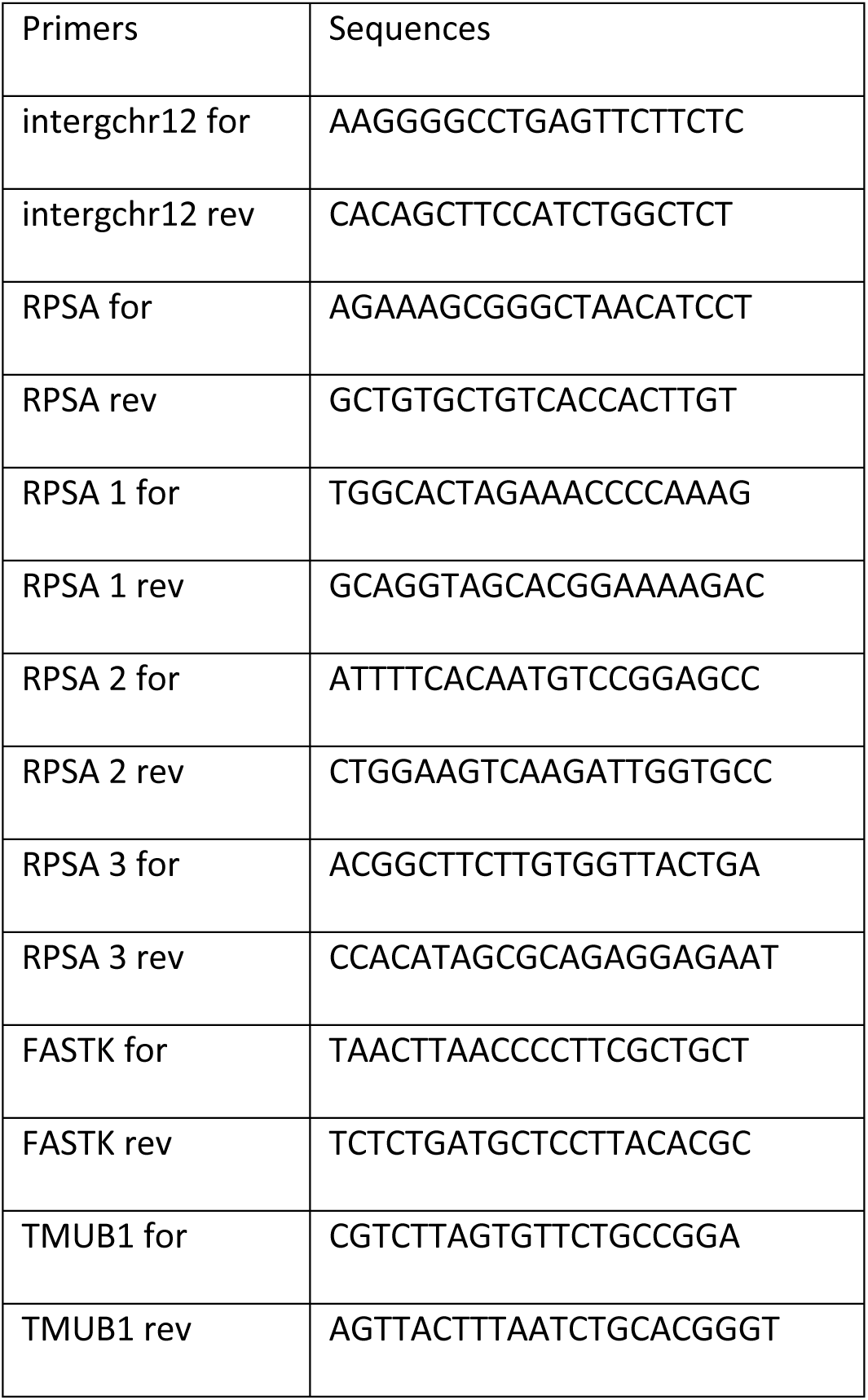
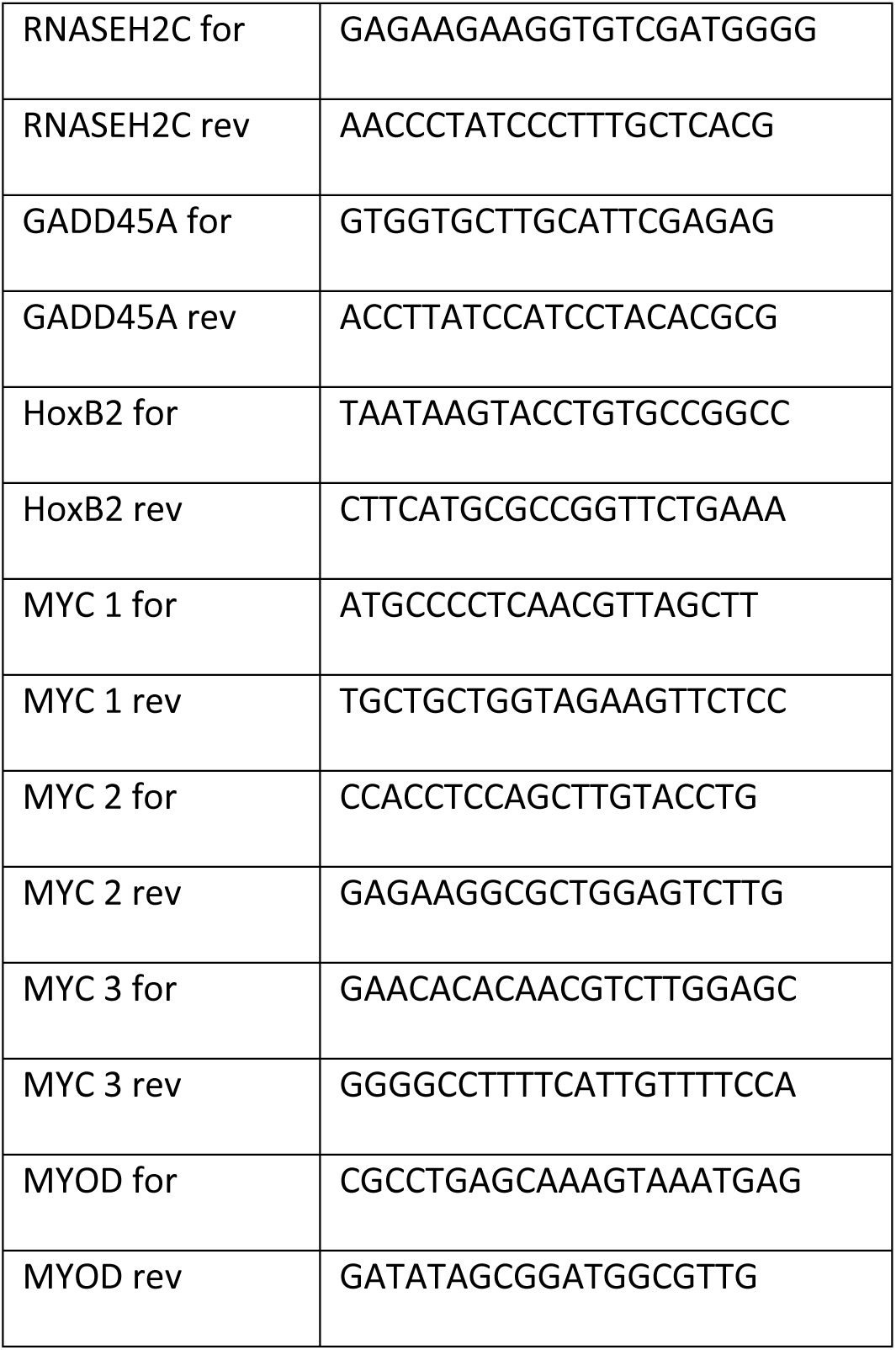

#### CRISPR/Cas9-mediated KO of ZMYND11 in K562 cells

K562 CRISPR KO cell lines were established as described in (Agudelo, Duringer et al. 2017) using two gRNAs to remove a full exon in C-term of ZMYND11 in AAVS1 cell lines expressing MBTD1 WT or truncated ZMYND11 or the ZMYND11-MBTD1 fusion. MBTD1 WT AAVS1 cell line is used as control. Target sequence for ZMYND11 (5’-ACACGCCGTCGTTTGTGGTCTGG-3’) was built in eSpCas9 pX330 backbone and (5’-ATTAACGGGCATCATACCTGTGG-3’) was built in MLM3636 backbone. For targeting with the CRISPR/Cas9 system, 4.10^5^ cells were transfected with 3 μg of each gRNA plasmid by Lipofectamine 3000 (Thermo Fisher). Three days post-transfection, cells were selected with ouabain. 5 days post-selection, cells were diluted for colony isolation and amplification followed by PCR screening. KO clones were validated by PCR screening, sequencing of the alleles, and RT-qPCR. *36B4* locus is used as control.

RNA extraction was performed with Monarch Total RNA Miniprep Kit (NEB #T2010) following the manufacturer’s indications. Then, 1µg of RNA was reverse transcribed by oligo-dT and random priming into cDNA with a qScript cDNA SuperMix kit (QuantaBio-VWR), according to the manufacturer’s instructions. Quantification of the amount of cDNA was done with SYBR Green I (Roche) on a LightCycler 480 (Roche) for real-time PCR,

RNA samples from 2 different validated KO clones were compared to 2 WT control. The following primers are used to validate KO ZMYND11 clones by PCR and RT-qPCR: PCR for (5’-ATTATGTGGCTTCGTTTGGC-3’); PCR rev (5’-TGAGCAGTACACAGTAATATGGG-3’); RT-qPCR RPLPO for (5’-CGACCTGGAAGTCCAACTAC-3’); RPLPO rev (5’-ATCTGCTGCATCTGCTTG-3’); ZMYND11 N-term for (5’-ATGTCTCGAGTCCACGGTATG-3’); ZMYND11Nterm rev (5’-GCCCATCTCCTGTTTGTTTGT-3’); ZMYND11 C-term for (5’-CCAAGAATGCTGCATCGGAG-3’); ZMYND11 C-term rev (5’-CTTCCATTTCAGAACGCAGCT-3’).

#### Library preparation and long-read RNA-sequencing

Cell lysates were homogenized with QIAshredder columns (QIAGEN #79654) and total RNA was extracted with RNeasy mini columns (QIAGEN #74104). RNA was DNase treated using TURBO DNase (ThermoFisher Scientific #AM2238). RNA quality was verified on a 2100 Bioanalyzer (Agilent Technologies) and polyadenylated RNA was enriched using NEBNext Poly(A) mRNA Magnetic Isolation Module (NEB # E7490L). Libraries were generated using 100 ng of poly(A)+ RNA and Oxford Nanopore Technologies’ Direct cDNA Sequencing Kit (ONT SQK-DCS109), with each sample barcoded with the Native Barcoding Expansion (ONT EXP-NBD104), according to the manufacturer’s specifications. Barcoded samples were pooled, loaded on a MinION sequencer using four R9 Flow Cells (ONT FLO-MIN106D), and sequenced using the MinKNOW software v19.

#### Long-read RNA-seq data analysis

Raw fast5 files were basecalled using Guppy v.4.2.2 with the guppy_basecaller command and the dna_r9.4.1_450bps_fast.cfg configuration file and default settings. Output fast5 files were generated with the –fast5_out argument. Barcodes were detected and reads were separated with the guppy_barcoder using the default settings. The resulting FASTQ reads were aligned to the GRCh38 human reference with Minimap2 using the following parameters: -aLx splice --cs=long (Li 2018). Raw read counts were obtained with the featureCounts tool from the Subread package v 2.0.0, using the exon counting mode (Liao, Smyth et al. 2014).

EdgeR R-package (v3.12.1) was then used to normalize the data, calculate RNA abundance at the gene and transcript levels (as counts per million reads (CPM), and perform statistical analysis (Robinson, McCarthy et al. 2010). Briefly, a common biological coefficient of variation (BCV) and dispersion (variance) were estimated based on a negative binomial distribution model. This estimated dispersion value was incorporated into the final EdgeR analysis for differential gene expression, and the generalized linear model (GLM) likelihood ratio test was used for statistics, as described in EdgeR user guide. Differential gene expression was established as genes with p-value ≤0.01 and FDR ≤0.25 and overlaps between comparisons were assessed using InteractiVenn (Heberle, Meirelles et al. 2015).

#### Gene ontology (GO) term enrichment analysis

GO term enrichment was assessed using the Database for Annotation, Visualization and Integrated Discovery (DAVID 6.8) (Jiao, Sherman et al. 2012). Biological processes (BP), cellular components (CC) and pathway enrichment with Kyoto Encyclopedia of Genes and Genomes (KEGG) annotations were analyzed. An FDR ≤ 0.1 was set as cut-off for significant enrichment.

#### Integration of long-read RNA-seq and ChIP-seq datasets

RNA-seq data was compared to previously established ChIP-seq binding of the MBTD1-ZMYND11 fusion. Genes were binned as up- or down-regulated by the fusion protein, as well as bound or not bound by the fusion protein. Non-random association between these two categorical variables was assessed using the Fisher’s exact test.

#### Analysis of alternative splicing

Analysis of variations in splice isoform expression was performed by comparing CPM values and differential expression obtained from EdgeR at the gene and transcript levels. For the previously identified genes showing differential expression between at least two samples (p-value ≤0.01 and FDR ≤0.25), those with differentially expressed transcripts as well (p-value ≤0.01 and FDR ≤0.10) were selected. Gene expression (in Z-scores of Log_2_(CPM)) was clustered hierarchically using the Ward.D2 clustering method and Euclidean clustering distance, and graphed alongside corresponding transcript expression, using the ComplexHeatmaps R package (Gu, Eils et al. 2016). Heatmaps were next separated to show the genes and transcripts differentially expressed in the same, or in opposite directions.

Differentially expressed transcripts (p-value ≤0.01 and FDR ≤0.10) corresponding to genes that showed no differential expression (p-value ≥0.01 and FDR ≥0.25) were then selected. To highlight the genes with high changes in transcript expression compared to gene expression, these selected genes were subjected to an additional cut-off of ≥2-fold difference in Log2(Fold change) values at the transcript and gene levels, between the selected sample and control. Expression for the resulting list of transcripts was graphed, and overlap between transcripts differentially expressed between each sample comparisons were presented as Venn diagrams using InteractiVenn (Heberle, Meirelles et al. 2015).

Transcriptome-wide evaluation of alternative splicing events was performed with PSI-sigma v1.9 (Lin *et al*., Bioinformatics, 2019, doi:10.1093/bioinformatics/btz438). Long-read RNAseq samples were compared one-on-one using the following parameters : -nread 10 --type 2 --fmode 3. Results were filtered by applying cut-offs for at least 10% variation in percent spliced-in values between samples (ΔPSI), p-value ≤ 0.05 and false discovery rate (FDR) ≤ 10%. Significant alternative 5’ and 3’ splice site events were grouped together as “Alternative splice site”, while single and multiple exon skipping events were grouped as “Exon skipping”. Significant divergence of observed splicing event counts, compared to expected counts, was determined via Chi-square test of independence with a level of significance of 0.05.

#### Reverse Transcription-qPCR

In ZMYND11 KO K562 cells stably expressing ZMYND11-MBTD1 fusion, MBTD1, WT ZMYND11, truncated ZMYND11 or empty vector, total RNA was extracted with Monarch Total RNA Miniprep Kit. 1ug of RNA was reverse transcribed by oligo-dT and random priming into cDNA with a qScript cDNA SuperMix kit (QuantaBio-VWR), according to the manufacturer’s instructions.

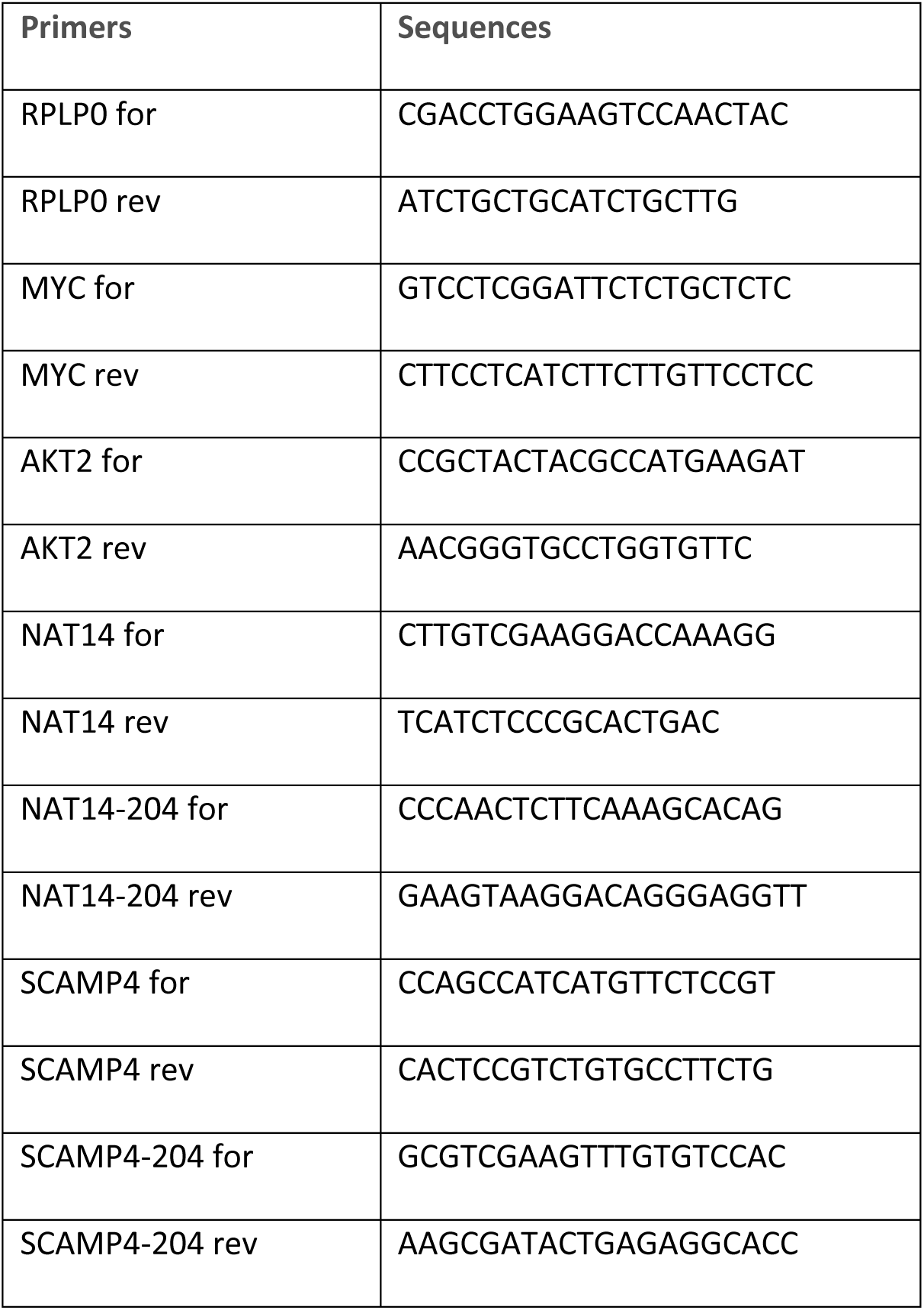

#### Overexpression of ZMYND11 and MBTD1 constructs during mouse ESC differentiation

For transient transfections, the same constructs as for the generation of the K562 isogenic cell lines were used. R1 ESCs were dissociated using TrypLE^TM^ Express enzyme 1X (ThermoFisher Scientific #12604021) and seeded on 0.1% gelatin in 2i-LIF medium at 125,000 cells/well in 6-well plates. The following day, the cells were transfected with the different constructs using Lipofectamine^TM^ Stem Transfection Reagent (ThermoFisher Scientific #STEM00001) following the manufacturer’s instructions, with 2 µg of DNA and 5 µL of lipofectamine/well. Cells were selected for 2 days with puromycin (1 μg/mL) in FBS-LIF medium (15% FBS plus LIF at 20 ng/mL, 1 mM sodium pyruvate, 2 mM L-glutamine, and β-mercaptoethanol (10^-4^ M) in DMEM high-glucose) and then seeded on 0.1% gelatin in N2B27 medium (140,000 cells/dish in 6cm-dishes) for differentiation. Medium was changed 2 days later and cells were harvested 4 days post seeding in N2B27 medium.

For stable transfections, the different constructs were cloned in a piggyBac transposon (PB) destination vector containing a doxycycline-inducible TetO promoter, a Gateway cloning cassette as well as the constitutive rtTA and puromycin resistance genes (TetO-CMV-gateway-IRES-mCherry-rtTA-2A-puromycin vector) by Gateway reaction (ThermoFisher Scientific #11789013 #11791020) following the manufacturer’s instructions. Cells were transfected as described previously and selected for 3 days with puromycin (1 µg/mL) in 2i-LIF medium. For differentiation, the obtained cell lines were seeded on Geltrex™ LDEV-Free, hESC-Qualified, Reduced Growth Factor Basement Membrane Matrix (Thermo Fisher Scientific #A1413302) in 2i-LIF medium (350,000 cells/dish in 6cm-dishes). The next day (D0), medium was replaced with EpiLC medium (20 ng/mL rhActivin A (PeproTech #120-14E), 12 ng/mL rhFGF2 (PeproTech #100-18B), and 1% KSR (ThermoFisher Scientific #10828028) in N2B27 medium) with addition of 1 μg/mL doxycycline (Sigma-Aldrich #D9891-1G) to induce the expression of the transgenes. Cells were split with StemPro^TM^ Accutase^TM^ Cell Dissociation Reagent (ThermoFisher Scientific #A1110501) after two days and seeded again at 350,000 cells/dish in the aforementioned media and matrix. Cells were harvested at D0, D2 and D5.

For both transient and stable transfections, total RNA was extracted using the EZ-10 Total RNA Miniprep Kit (BioBasic #BS88583) and DNase I (RNAse-free, 1U/µL)-treated (ThermoFisher Scientific #EN0521) following the manufacturers’ instructions. RNA was reverse-transcribed for RT-qPCR using iScript^TM^ gDNA Clear cDNA Synthesis Kit (Bio-Rad #1725035) following the manufacturer’s instructions. qPCR was performed using SsoAdvanced Universal SYBR Green Supermix (Bio-Rad #1725274). The mean relative fold-change in mRNA expression was normalized to *Rps13* housekeeping gene expression in the control condition (empty vector). The error bars represent the Standard Deviation (STD). Refer to table below for primer sequences.

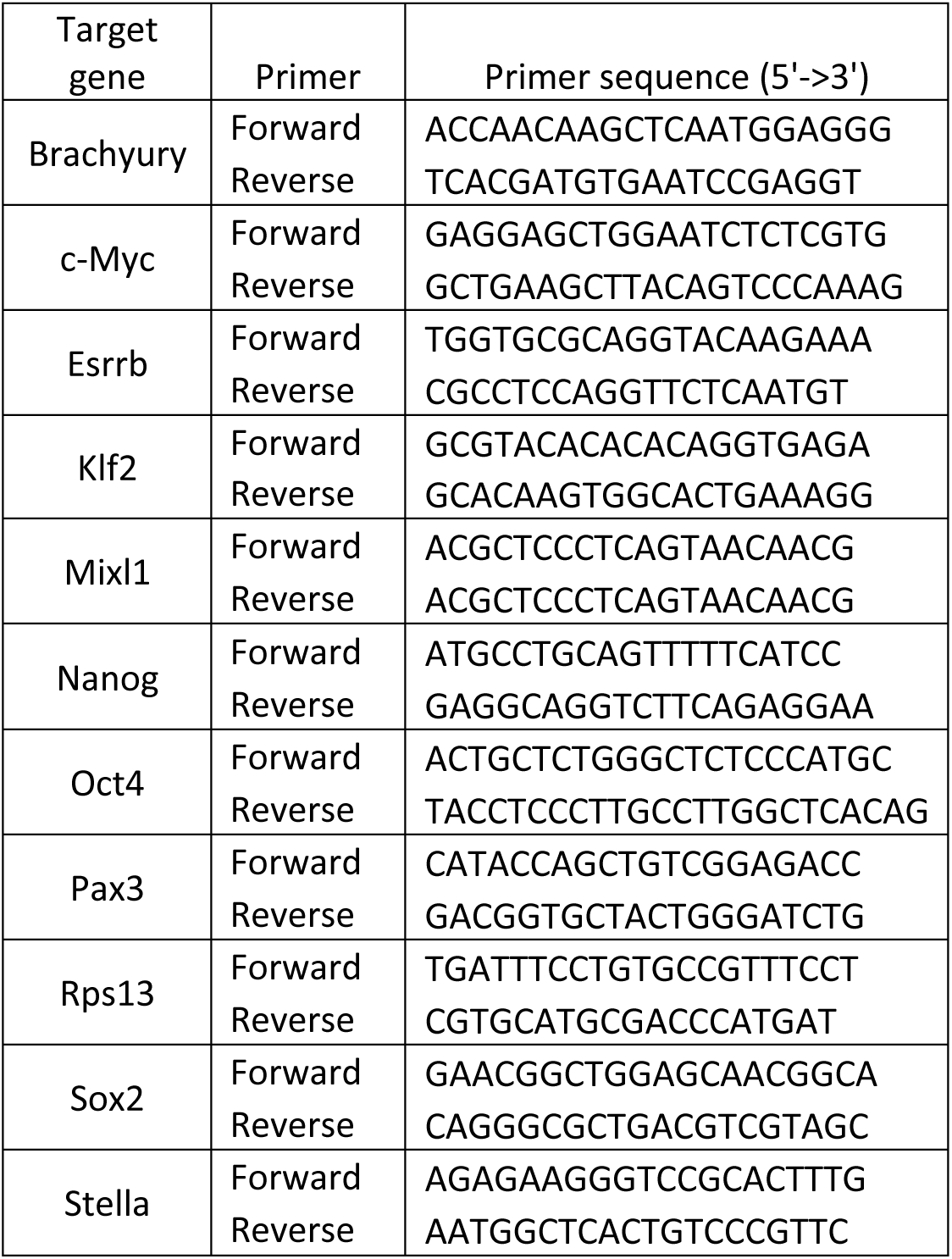

#### Colony-forming unit (CFU) assay

Bone marrow mononuclear cells harvested from 8-12 week old C57BL/6 mice were lineage depleted with EasySep™ Mouse Hematopoietic Progenitor Cell Isolation Kit (STEMCELL Technologies #19856) following manufacturer protocol. Mouse Lin^-^ cells were seeded at a density of 100,000 cells/well in a 24-well plate and cultured in SFEM II media (STEMCELL Technologies #09655) supplemented with 10% FBS, mSCF (100ng/mL, R&D Systems #455-MC-010), mTPO (100ng/mL, PeproTech #AF-315-14), mIL3 (10ng/mL, R&D Systems #403-ML-050), mIL6 (10ng/mL, R&D Systems #406-ML-025), and 1X pen/strep (ThermoFisher Scientific #15140122). C57BL/6 Lin-cells were infected with lentivirus for 72 hours at an MOI of 50. Transduced cells were isolated by tagBFP marker expression through FACS (BD FACS ARIA Fusion), resuspended at 8,000 cells/mL in ColonyGEL Mouse Complete Medium (ReachBio #1202), and plated at 1mL/plate in 35mm dishes (Corning #430588). Plates were imaged with a STEMvision™ instrument (STEMCELL Technologies #22005) after 7 days of incubation, and colonies were manually counted.

#### Cloning of lentiviral overexpression constructs into pSMALB

Lentiviral constructs were made by subcloning cDNAs from the same vector used in mESCs into the lentiviral vector pSMALB (a gift from John Dick & Peter van Galen (van Galen, Kreso et al. 2014)), downstream of an SFFV promoter, using Gateway LR Clonase (Thermofisher Scientific #11791020) following manufacturer instructions.

#### Lentiviral Production

Lenti-X cells (Takara Bio #632180) were co-transfected with 23ug lentiviral overexpression vector, and 9ug pMD2.G and 13ug psPAX2 packaging plasmids (Addgene) per T150 flask (Corning #CLS430825), using 92ul Lipofectamine 2000 (ThermoFisher Scientific #11668019). Media was changed 8 hours post transfection and viral supernatant was collected after 72 hours. The supernatant was centrifuged and filtered through 0.45µm Steriflip-HV filters (Millipore #SE1M003M00) before concentration by ultracentrifugation (20,000 RPM for 2.5 hours, Optima L-100XP unit with 70Ti rotor, Beckman Coulter). The viral particle pellet was re-suspended in SFEM II (STEMCELL Technologies #09655) and stored at -80 °C.

#### Recruitment-activator assay

This assay is described in (Alerasool et al. 2022). The reporter HEK293T cell line with a TRE3G-EGFP reporter was a kind gift from Stanley Qi (Stanford University). The reporter cell line was transduced with a lentivirus expressing ABI-dCas9 followed by two rounds of selection under 6µg/ml blasticidin. These cells were then transduced with a lentivirus expressing EBFP2 and a gRNA targeting seven tetO repeats in the TRE3G promoter (gRNA sequence GTACGTTCTCTATCACTGATA). Single-cell derived clonal cell lines were generated and a clone showing robust EGFP induction by a strong transcriptional activator VPR was selected for downstream assays. 96-well plates were seeded with 3x10^4^ cells per well one day prior to transfection. 150 ng of each construct was transfected using polyethylemine (PEI). Transfected cells were induced 24 hours after transfection by treatment with 100 µM abscisic acid. 48 hours after induction, cells were dissociated and resuspended in flow buffer using a liquid handing robot and analyzed by LSRFortessa (BD). Flow cytometry data was analyzed using FlowJo by gating for positive gRNA (EBFP2), then further for construct (TagRFP) expression. At least 25,000 cells were analyzed for each replicate.

### QUANTIFICATION AND STATISTICAL ANALYSIS

A mock cell line 3xFlag-2xStrep is used as controls. ChIP-qPCRs to validate ChIP-seq and RT-qPCRs to validate RNA-sequencing were performed in biological duplicate. Error bars represent the range from two independent replicates. Statistical analyses were performed via two-way ANOVA using Prism version 7 (GraphPad software inc. California, USA) followed by Tukey’s test unless otherwise noted. P-values <0.05 were considered significant.

## SUPPLEMENTARY FIGURE LEGENDS

**Figure S1.**
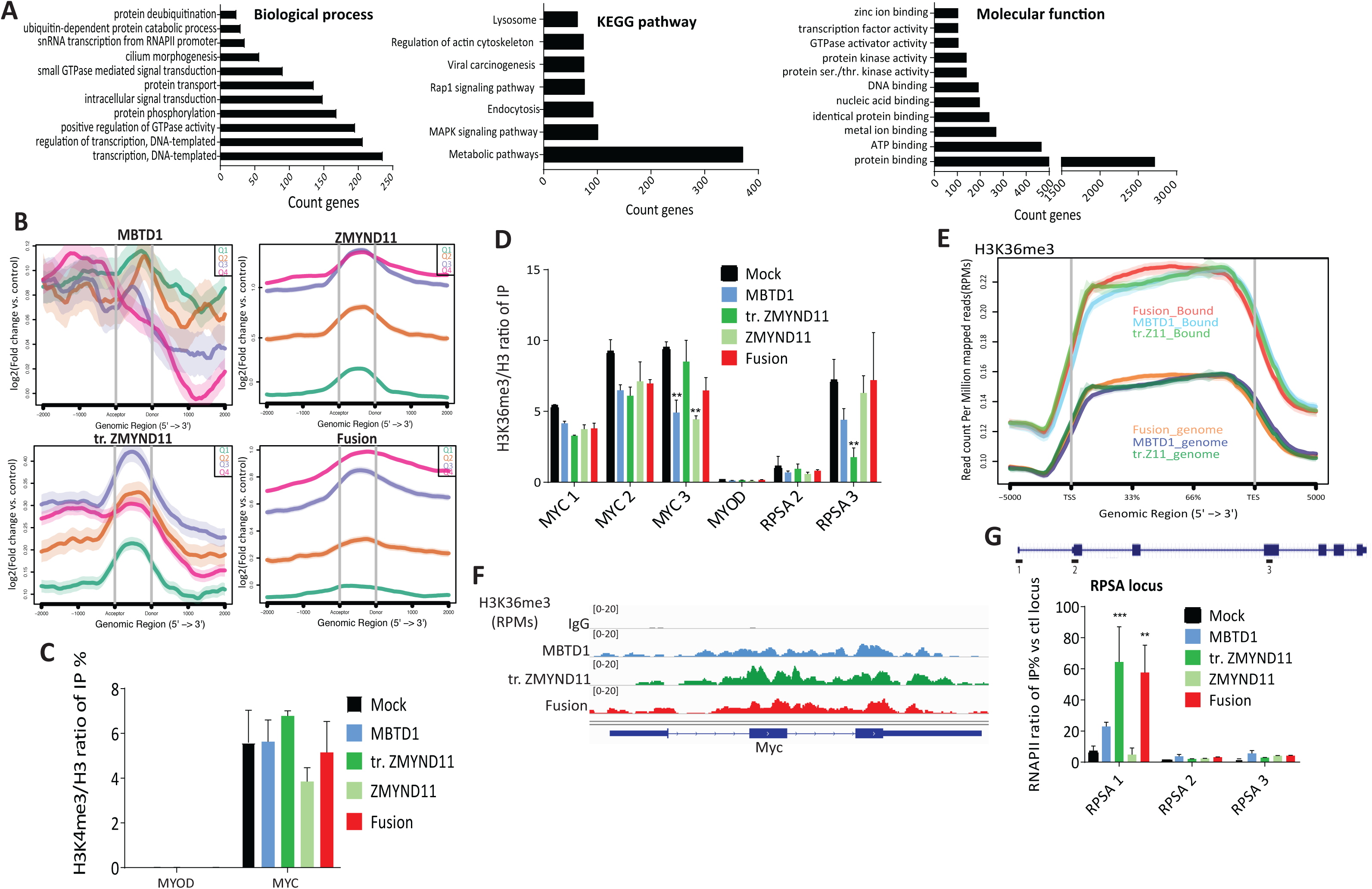
The ZMYND11-MBTD1 fusion is enriched in gene bodies, related to figureA. Gene Ontology (GO) term enrichment analysis using DAVID 6.8 shows terms for biological processes, molecular function and KEGG pathway (FDR ≤ 0.01) enriched in ZMYND11-MBTD1 fusion and truncated ZMYND11 but not MBTD1. **B.** Average profile of ChIP peaks binding to acceptor and donor splice sites. Genes are separated by gene expression quantiles (Q4 is the highest expression). MBTD1 is not enriched on these genomic regions compared to the three others, ZMYND11, tr. (truncated) ZMYND11, and the fusion. **C.** ChIP-qPCR of H3K4me3 in K562 cells stably expressing MBTD1, WT ZMYND11, truncated ZMYND11 or the ZMYND11-MBTD1 fusion. Signals are presented as a ratio of IP % on total H3 to correct for nucleosome occupancy at *MYC* locus and *MYOD* locus as control. **D.** ChIP-qPCR of H3K36me3 from cells in C along *MYC* and *RPSA* loci (see Fig. 2F and G below for positions). *MYOD* locus is used as negative control. Signals are presented as a ratio of IP % on total H3 to correct for nucleosome occupancy. **E.** Metagene analysis showing anti-H3K36me3 CUT&RUN-seq signals across genes specifically bound (Bound) or not (genome) by the fusion or its partners, in K562 cells stably expressing the ZMYND11-MBTD1, MBTD1 or tr. (truncated) ZMYND11. **F.** Genome browser track of the *MYC* locus from anti-H3K36me3 CUT&RUN-seq in K562 cells expressing the fusion or its partners MBTD1 and tr. (truncated) ZMYND11. Signals are indicated are reads per million reads (RPMs). IgG track is shown as background control.**G.** ChIP-qPCR of RNA polymerase II (RNAPII) signals are presented as a ratio of IP % on negative locus (*MYOD*). The set of primers used are shown in the schematic *RPSA* locus. Error bars represent the range based on two independent experiments. Statistical analyses were performed by two-way ANOVA test followed by Tukey’s test, *, p < 0.05 , **, p < 0.01, ***, p < 0.001, ****, p <0.0001.

**Figure S2:**
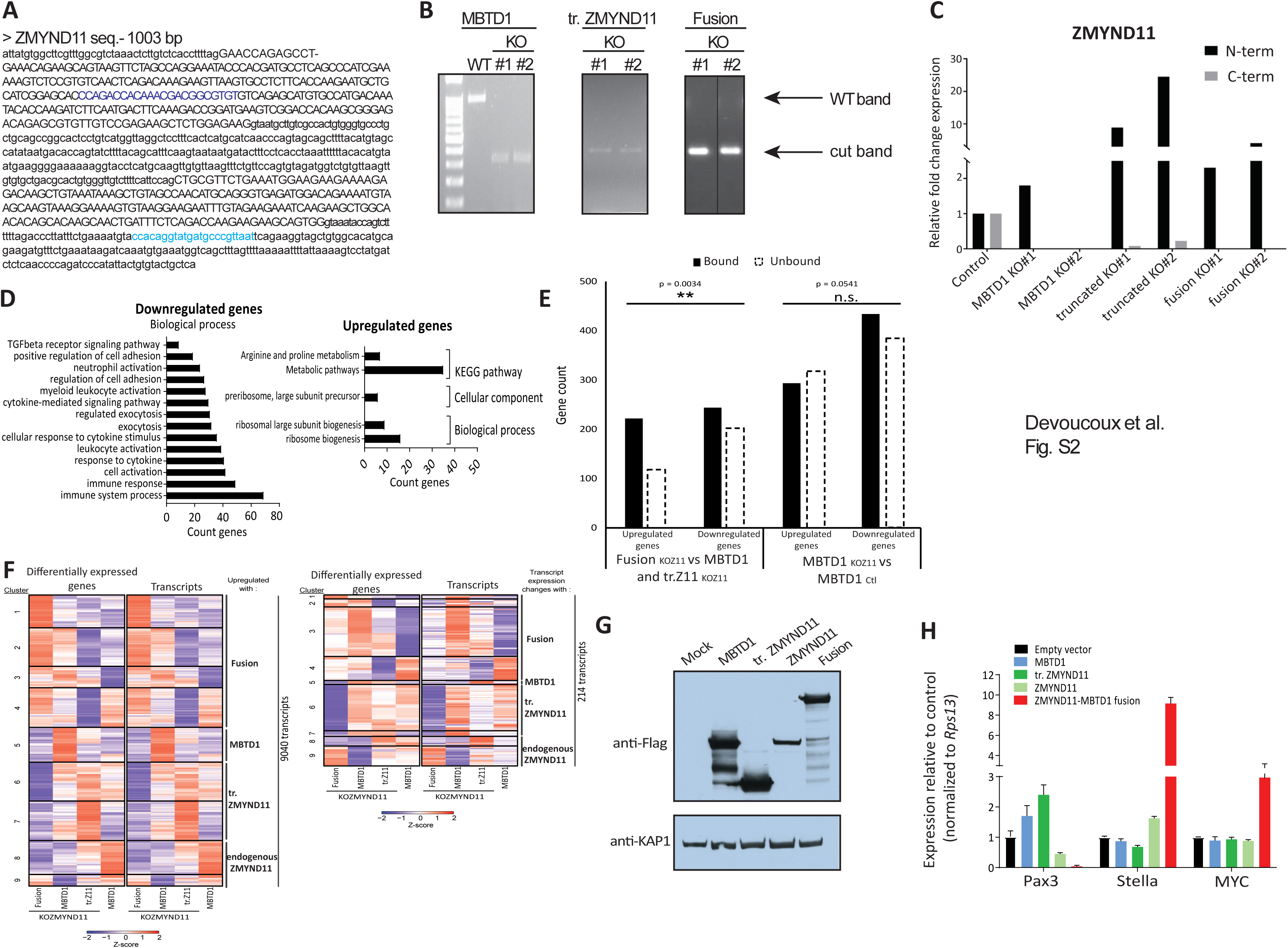
Oncogenic mechanisms due to the expression of ZMYND11-MBTD1 fusion, related to figures 3 and 4. **A.** *ZMYND11* DNA sequence showing both gRNAs used to generate a truncation in the MYND domain of ZMYND11 (in purple and in blue). **B.** Selection of *ZMYND11* knock-out (KO) clones. Results of a PCR-based screening to detect targeted deletion in single-cell-derived K562 clones after ouabain selection. Two homozygote KO clones are shown in each AAVS1 cell line (MBTD1, truncated (tr.) ZMYND11 and ZMYND11-MBTD1 fusion (fusion)). Alleles were also sequenced to verify the accurate deletion in each KO clone. **C.** RT-qPCR of ZMYND11 expression after endogenous *ZMYND11* KO. 2 sets of primers were used in N-term and C-term regions of the *ZMYND11* locus. Signals are presented as a fold expression relative to the *RPLP0* locus control. Cell lines expressing tr.ZMYND11 and the fusion from the *AAVS1* locus show signals from the N-term region because of the transgene. **D.** Gene ontology (GO) analysis for upregulated and downregulated genes using string-db software (FDR < 0.005). **E.** Histogram showing number of genes up- or downregulated upon expression of the ZMYND11-MBTD1 fusion protein compared to either tr. (truncated) ZMYND11 or MBTD1 in KO ZMYND11 cell lines, or in control cell lines with endogenous ZMYND11 compared to MBTD1 in KO ZMYND11 cell lines. Genes are classified according to whether they are bound by the fusion or not (from ChIP-seq). Non-random association between fusion binding and up-or downregulation in RNA-seq samples was tested for using the Fisher’s exact test (**: p-value ≤ 0.01). Binding of the fusion is specifically linked to upregulation of genes. **F.** Heatmap of expression (in Z-scores of Log_2_(CPM)) for differentially expressed genes and their associated transcripts in genes with homodirectional changes in the gene and transcript expression (left), or in genes with opposite changes in the gene and transcript expression (right) in the indicated samples. Total numbers of transcripts observed are indicated. **G.** Western blot analysis of ZMYND11-MBTD1 (fusion) overexpression in mouse embryonic stem cells (mESCs). MBTD1, truncated ZMYND11 (tr. ZMYND11), ZMYND11 and empty vector are used as controls. mESCs were transiently transfected, selected for 48h with puromycin before initiating spontaneous differentiation. **H.** RT-qPCR of indicated pluripotency (*Stella*, *MYC*) and differentiation (*PAX3*) markers after overexpression of the ZMYND11-MBTD1 fusion in mESCs. The experiments were done in duplicates. MBTD1, ZMYND11, truncated (tr.) ZMYND11 and empty vector were used as controls.

**Supplemental Table S1:**
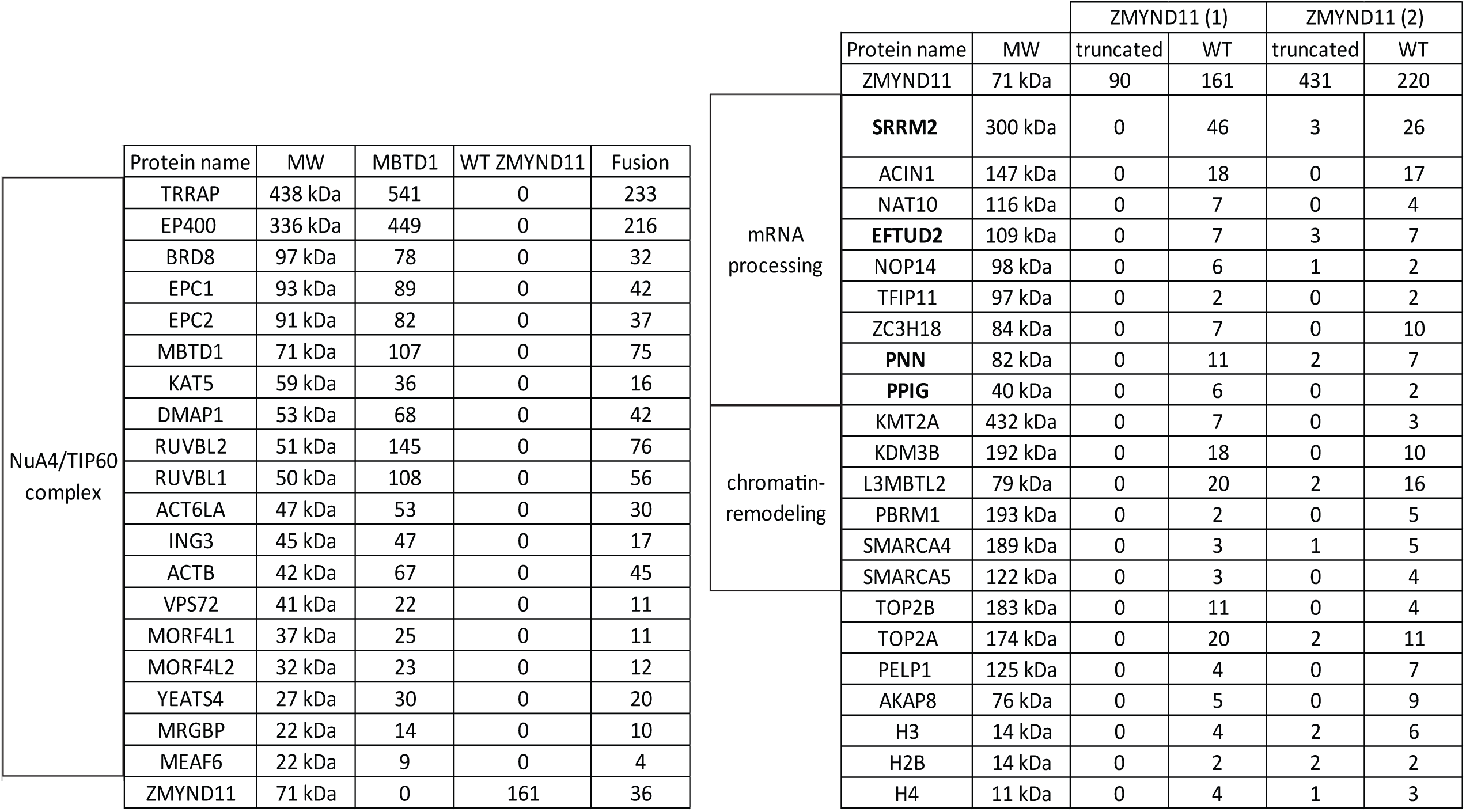
NuA4/TIP60 complex subunits identified (left) and ZMYND11 interactome (right) by Mass spectrometry analysis. Predicted molecular weights (MW) and total spectral counts for identified proteins are indicated. MBTD1, WT ZMYND11, truncated ZMYND11 and fusion indicate whether the subunits were expressed ectopically from the AAVS1 locus in K562 cells. In bold, ZMYND11 interactors previously identified by Guo et al, Mol cell, 2014. Related to figure 1.

**Supplemental Table S2:**
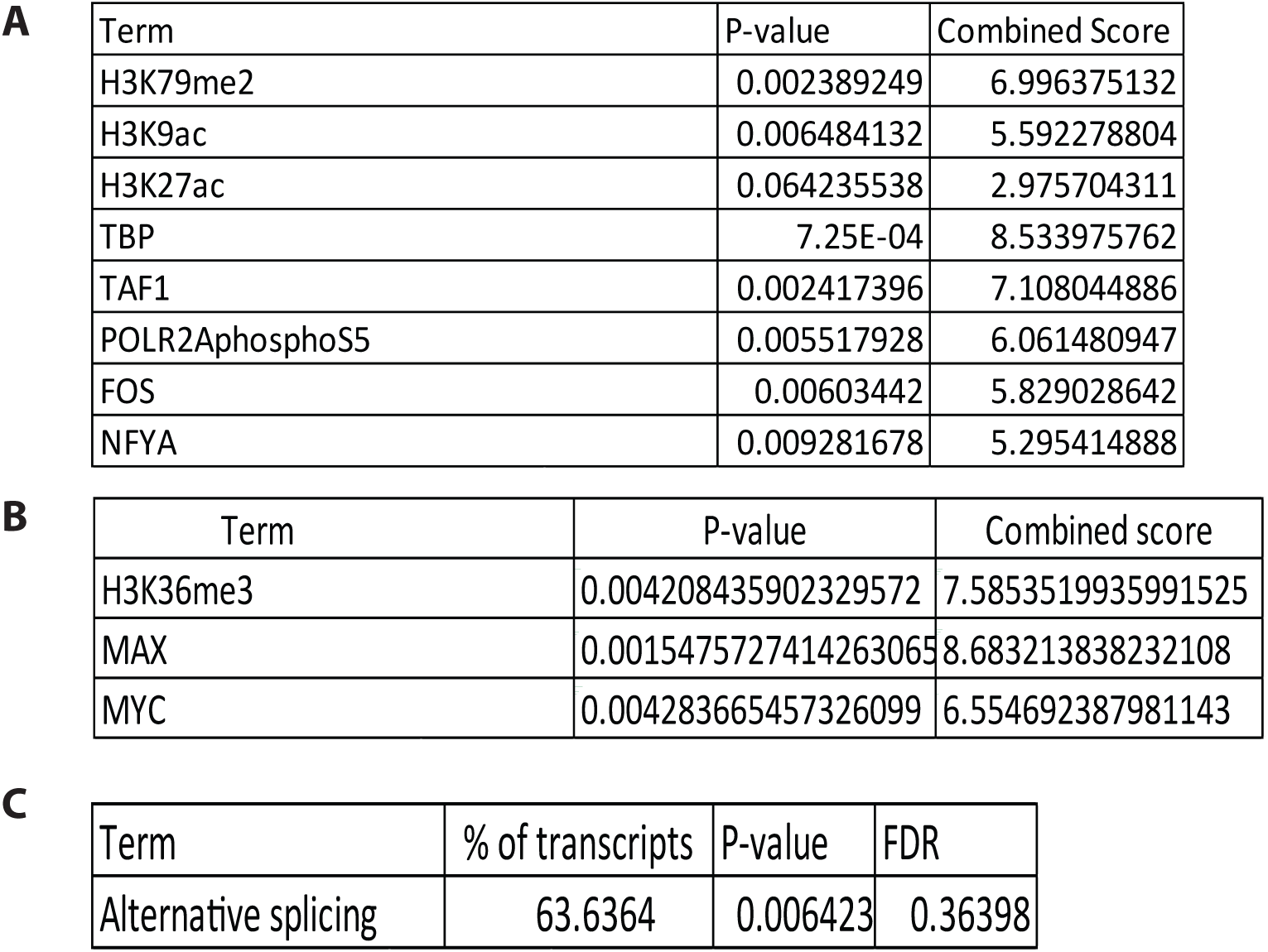
A. ChIP-sequencing analysis of the ZMYND11-MBTD1 fusion in K562: histone marks and transcription factors enriched on genes that are bound by ZMYND11-MBTD1 fusion protein and truncated ZMYND11 but not MBTD1. B. Long reads MiniON/expression analysis of genes upregulated by the fusion. (ENCODE Histone modifications 2015-EnrichR and ENCODE Transcription Factors ChIP-seq 2015-EnrichR). C. Functional analysis by DAVID 6.8 of genes linked to trancripts isoforms found affected by the ZMYND11-MBTD1 fusion. Related to figures 2, 3 and 4.

## Notes

### Competing Interest Statement

The authors have declared no competing interest.

